# Exploring the conformational landscape of a lanthipeptide synthetase using native mass spectrometry

**DOI:** 10.1101/2020.09.01.277863

**Authors:** Nuwani W. Weerasinghe, Yeganeh Habibi, Kevin A. Uggowitzer, Christopher J. Thibodeaux

**Affiliations:** Department of Chemistry & Centre de Recherche en Biologie Structurale, McGill University, 801 Sherbrooke St. West Montréal, Québec, Canada, H3A 0B8

## Abstract

Lanthipeptides are ribosomally-synthesized and post-translationally modified peptide (RiPP) natural products that are biosynthesized in a multistep maturation process by enzymes (lanthipeptide synthetases) that possess relaxed substrate specificity. Recent evidence has suggested that some lanthipeptide synthetases are structurally dynamic enzymes that are allosterically activated by precursor peptide binding, and that conformational sampling of the enzyme-peptide complex may play an important role in defining the efficiency and sequence of biosynthetic events. These “biophysical” processes, while critical for defining the activity and function of the synthetase, remain very challenging to study with existing methodologies. Herein, we show that native nanoelectrospray ionization coupled to ion mobility mass spectrometry (nanoESI-IM-MS) provides a powerful and sensitive means for investigating the conformational landscapes and intermolecular interactions of lanthipeptide synthetases. Namely, we demonstrate that the class II lanthipeptide synthetase (HalM2) and its non-covalent complex with the cognate HalA2 precursor peptide can be delivered into the gas phase in a manner that preserves native structures and intermolecular enzyme-peptide contacts. Moreover, gas phase ion mobility studies of the natively-folded ions demonstrate that peptide binding and mutations to dynamic structural elements of HalM2 alter the conformational landscape of the enzyme, and that the precursor peptide itself exhibits higher order structure in the mass spectrometer. Cumulatively, these data support previous claims that lanthipeptide synthetases are structurally dynamic enzymes that undergo functionally relevant conformational changes in response to precursor peptide binding. This work establishes nanoESI-IM-MS as a versatile approach for unraveling the relationships between protein structure and biochemical function in RiPP biosynthetic systems.

## Introduction

Ribosomally-synthesized and post-translationally modified peptides (RiPPs) are a structurally and functionally diverse group of peptide natural products that share common biosynthetic characteristics (Figure 1A).^1^ All RiPPs are derived from genetically encoded precursor peptides that are post-translationally modified by biosynthetic enzymes. Following post-translational modification, the modular RiPP precursor peptide is typically proteolyzed to separate the *N*-terminal leader peptide (which confers biosynthetic enzyme binding) from the *C*-terminal core peptide which contains the post-translationally modified amino acid residues. The modified core peptide is then exported from the cell where it exerts a biological function. RiPP biosynthetic pathways have recently attracted much interest from the natural products community for a number of reasons. First, RiPP natural products possess potent antimicrobial and other activities of biomedical relevance. Second, many RiPP biosynthetic enzymes catalyze the iterative, stepwise modification of their precursor peptides, and thus possess an inherently relaxed substrate specificity that may be useful for engineering novel biologically active peptides. Third, both RiPP biosynthetic enzymes and their precursor peptide substrates are genetically encoded – opening the door for the application of powerful molecular engineering approaches to generate and screen libraries of structurally complex peptides for novel biomedical functions.^2–5^ For all of these reasons, efforts to understand the mechanisms of substrate recognition and catalysis in RiPP biosynthetic enzymes are warranted.

**Figure 1.**
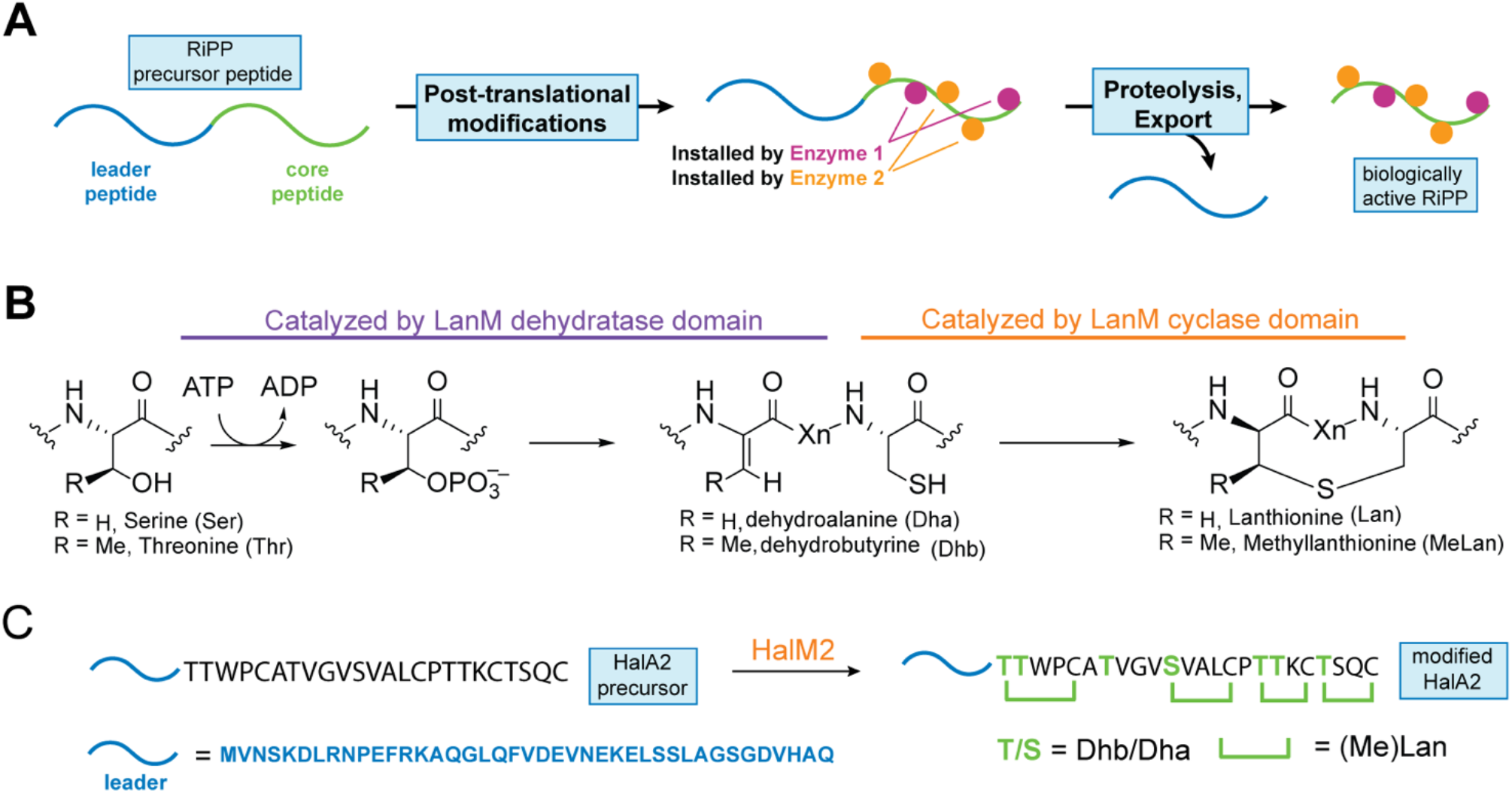
Overview of RiPP and Class II Lanthipeptide Biosynthesis. A) General features of RiPP biosynthesis, highlighted by the structurally modular, genetically encoded RiPP precursor peptide, the involvement of iteratively-active biosynthetic enzymes, and the proteolytic removal of the post-translationally modified core peptide. B) Chemical mechanism of thioether bridge formation catalyzed by class II lanthipeptide synthetases (LanM enzymes). The net dehydration and cyclization reactions are catalyzed in separate active sites of the enzyme. C) Post-translational modification of the HalA2 precursor peptide by the HalM2 synthetase, involving the dehydration of seven Ser/Thr residues and the formation of four thioether bridges.

While our understanding of the chemical mechanisms of RiPP biosynthetic enzymes have been rapidly expanding with the discovery and characterization of many new biosynthetic pathways, elucidation of the structural features that endow these enzymes with their relaxed substrate specificities and interesting catalytic properties have lagged behind. Nevertheless, high-resolution structures have been solved for a handful of enzymes involved in the biosynthesis of several types of RiPP natural products including the class I and II lanthipeptides,^6–10^ cyanobactins,^11–12^ orbitides,^13^ sactipeptides,^14^ lasso peptides,^15^ thiopeptides,^16^ and others.^17–19^ In several systems, co-crystal structures of the RiPP biosynthetic enzyme bound to its cognate precursor peptide have revealed the presence of a winged helix-turn-helix motif (termed the RiPP recognition element^20^) involved in leader peptide binding, but this RiPP binding element is not conserved in all RiPP biosynthetic enzymes, suggesting that other modes of peptide recognition exist. Despite these recent advances in RiPP structural biology, the mechanisms of core peptide recognition by RiPP enzyme active sites, how (or if) these interactions change during peptide maturation, and what role leader peptide binding might play in allosteric activation of the biosynthetic enzymes remain largely undefined. Clarification of these biophysical processes may ultimately inform rational approaches to engineer the catalytic properties and substrate specificities of RiPP biosynthetic enzymes.

Recently, we reported the use hydrogen-deuterium exchange mass spectrometry (HDX-MS) as a versatile biophysical tool for studying the structural dynamics of RiPP biosynthetic enzymes.^21^ Our studies focused on the well-characterized class II lanthipeptide synthetase (HalM2) involved in the biosynthesis of the *β* peptide of haloduracin (Figure 1 C), a potent two-component lanthipeptide antibiotic produced by *Bacillus halodurans*.^22^ Like all class II lanthipeptide synthetases, HalM2 is a bifunctional enzyme that uses separate active sites to catalyze the ATP-dependent dehydration of Ser/Thr residues in the precursor lanthipeptide (HalA2), followed by the Zn-dependent intramolecular Michael-type addition of Cys residues onto the nascent dehydration sites to make (methyl)lanthionine thioether rings (Figure 1B).^9, 23–25^ Despite the inherently low (i.e. peptide-level) spatial resolution of the bottom-up HDX-MS approach, these studies allowed us to identify a novel precursor peptide binding element in HalM2, as well as previously overlooked dynamic protein loops that exerted substantial perturbations to substrate binding and HalM2 catalytic activity when mutated. In addition to these mechanistic insights into lanthipeptide biosynthesis, the HDX-MS study highlighted the power of mass spectrometry-based techniques for revealing important functional information on a RiPP biosynthetic enzyme for which no high-resolution structure is available.

In the present study, we report a second emerging mass spectrometry-based technique for studying the structural properties of RiPP biosynthetic enzymes. Namely, we report to our knowledge the first detailed application of native electrospray ionization mass spectrometry to study the structural properties and biophysical interactions of a folded RiPP biosynthetic enzyme (HalM2) in the gas phase. The approach utilizes nanoelectrospray ionization (nanoESI) to allow for the rapid and gentle desolvation of protein ions from aqueous solvent without the need for excessive droplet heating or the application of strong electric fields, which can lead to protein denaturation. Using collisional cross section values calculated from gas phase ion mobility (IM) separations, we show that the tertiary structure of the HalM2 ions closely resembles the structure of a HalM2 homology model, strongly suggesting that the gas phase HalM2 ions possess a near-native conformation under nanoESI conditions. In support of this conclusion, we show that HalM2 remains non-covalently bound to its precursor peptide substrate (HalA2) in the gas phase, and that the binding affinity (*K*_*d*_) for HalA2 as measured from the MS ion signals closely matches the *K*_*d*_ value measured in solution. Moreover, we provide evidence that the HalA2 leader peptide and full-length precursor peptide also possess some degree of tertiary structure when electrosprayed from aqueous solvent droplets – allowing us to test whether RiPP precursor peptides are structured in solution. Finally, using IM measurements, we analyze the gas phase conformational landscapes of wt HalM2 and a series of mutant enzymes with altered structural dynamics and catalytic properties,^21^ in order to provide insight into the HalM2 structural elements that contribute to the enzyme’s fold and function. Thus, similar to the HDX-MS platform we recently reported,^21^ the application of native nanoESI coupled to ion mobility mass spectrometry (nanoESI-IM-MS) has provided a significant level of new insight into the structural properties of class II lanthipeptide synthetases and their precursor peptide substrates. Due to the sensitivity of these MS methods and their ability to probe both inter- and intramolecular interactions and conformational dynamics within complex mixtures of unlabelled biomolecules, mass spectrometry-based methods are poised to find great utility in the study of other structurally dynamic RiPP systems, providing access to information on functionally relevant conformational dynamics in these systems that are difficult to access by other methods.

## Results and Discussion

### Characterizing the enzymatic activity of HalM2 in native nanoESI solvent

To assess the feasibility of performing gas phase studies on natively-folded HalM2, we first exchanged HalM2 from its HEPES storage buffer into a solution of 200 mM ammonium acetate (pH 7.5) – a commonly used volatile solvent for native MS studies.^26^ When assayed for activity under otherwise standard reaction conditions,^27^ the enzyme exhibited indistinguishable catalytic properties in ammonium acetate (Figure S1), installing the expected four thioether rings into a seven-fold dehydrated HalA2 precursor peptide over the expected time scale.^22,27^ We next determined whether HalM2 maintained enzymatic activity in the nanoESI source. Accordingly, we prepared a reaction mixture in 200 mM ammonium acetate (pH 7.5) containing all of the components needed for HalM2 activity (HalA2 precursor peptide, ATP, MgCl_2_, and TCEP).^27^ The reaction mixture was loaded into a platinum-coated borosilicate nanoESI emitter, mounted to the nanospray ESI source of a Synapt G2-Si ion mobility mass spectrometer, and mass spectral data were recorded at 1 s intervals over 20 min. Under these conditions, the HalA2 precursor peptide yielded an intense signal that allowed us to follow the time-dependent, HalM2-catalyzed dehydration of the peptide in a continuous manner (Figure 2A). After approximately 16 min, the unmodified peptide starting material had been completely converted into the fully 7-fold dehydrated product. These data establish that HalM2 is fully active in the nanoESI source under the conditions used to perform native mass spectrometry. Moreover, the data illustrate the feasibility of continuous nanoESI-MS kinetic assays of RiPP biosynthetic enzymes, opening the door to more detailed mechanistic and kinetic characterization of these multifunctional catalysts.

**Figure 2:**
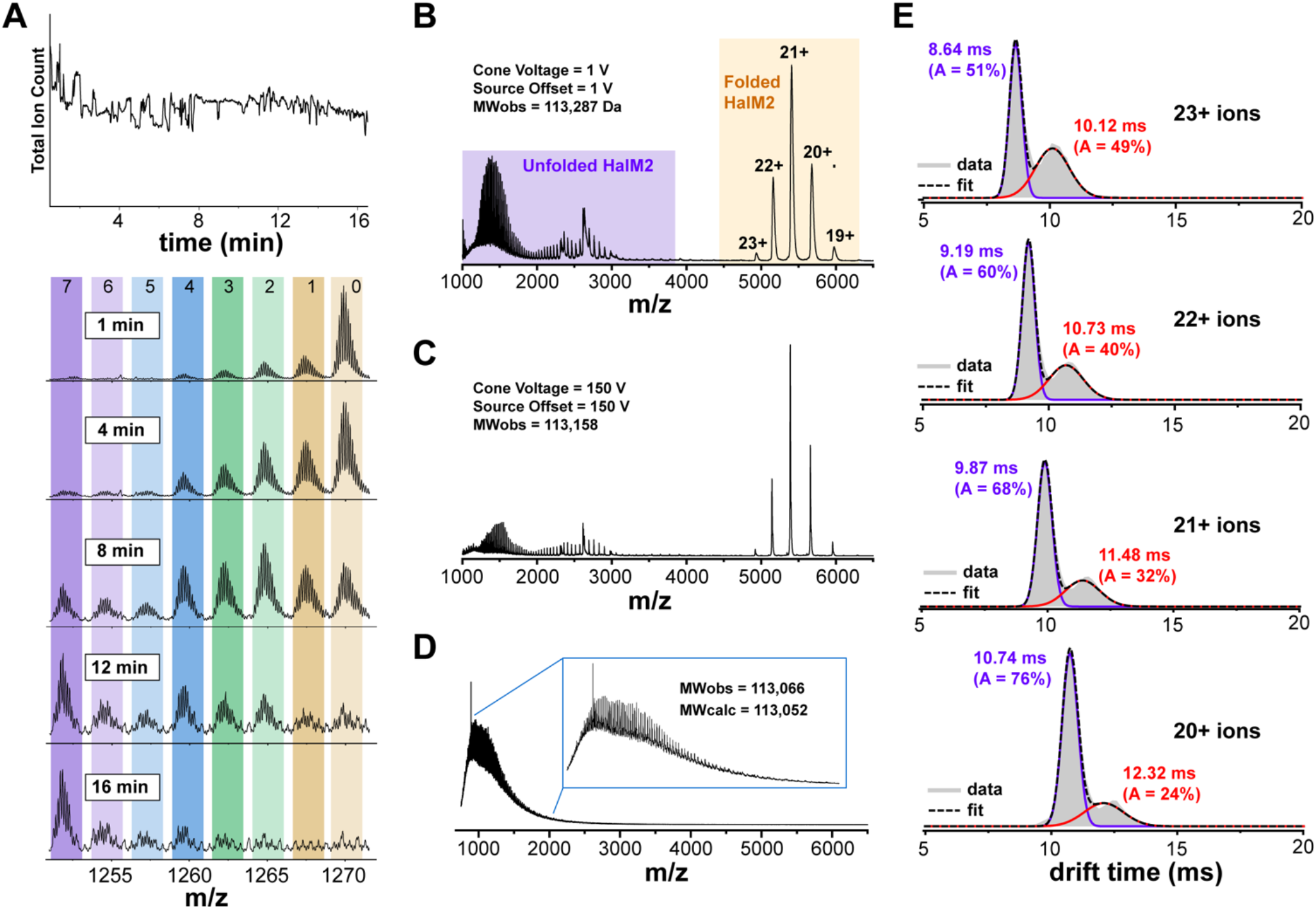
A) Continuous monitoring of HalM2 activity in the nanoESI source. A reaction mixture containing 0.5 μM HalM2, 50 μM HalA2, 1 mM ATP, 1 mM MgCl_2_, and 1 mM TCEP in 200 mM ammonium acetate (pH 7.5) was electrosprayed continuously over 16 min. The unmodified HalA2 starting material becomes progressively more dehydrated over time. The number of dehydrations (0 – 7) is indicated at the top of the time course. B) Native nanoESI-MS spectrum of 5 μM HalM2 collected with a capillary voltage = 1.5 kV, cone voltage = 1 V, a source offset = 1 V, and a source temperature of 35 °C. Folded HalM2 conformations are observable at higher m/z (lower charge states), while denatured HalM2 conformations are more extensively charged (lower m/z). C) Increasing the source voltages to 150 V enhances ion desolvation, narrows the peak width, and allows a more accurate determination of HalM2 molecular weight. D) HalM2 electrosprayed from 50% acetonitrile containing 0.1% formic acid is fully denatured and completely desolvated. E) Travelling wave ion mobility separation of the folded HalM2 20^+^ − 23^+^ ions was performed using a travelling wave velocity of 550 m/s and a wave height = 40 V at a nitrogen pressure of 3.2 mbar (see Table S5 for complete list of instrumental settings). The drift time distributions revealed the presence of two major HalM2 conformational populations at each charge state. From the measured drift times for each conformation (indicated above each peak), collisional cross section values (Ω_*c*_ and Ω_*0*_, Table S4) were calculated as described in the Supporting Information.

### Ion mobility mass spectrometry confirms a near native tertiary structure for gas phase HalM2 ions

Following these necessary control experiments, a 5 μM solution of HalM2 in 200 mM ammonium acetate was subjected to gentle nanoESI conditions (Table S5) in an attempt to maintain the enzyme in its native fold. Intense multiply charged ions corresponding to HalM2 charge states 19^+^ − 23^+^ were detectable in the m/z range 4500-6500 under these conditions, as well as more highly charged ions at lower m/z (Figure 2B). The 19^+^ − 23^+^charge states likely represent folded HalM2 conformations, where the majority of potential ionization sites are protected from charging during the ESI process. The molecular weight estimated from the 19^+^ − 23^+^ ions (*MW*_*obs*_ = 113,287 ± 25 *Da*, Table S1) is significantly larger than the expected HalM2 molecular weight (*MW*_*calc*_ = 113,052 *Da*), indicating incomplete desolvation of these ions under the gentle nanoESI conditions employed. A molecular weight closer to the expected value could be obtained by increasing the voltages in the nanoESI source (*MW*_*obs*_ = 113,158 *Da*, Figure 2C) or by fully denaturing the enzyme in an acetonitrile/water/formic acid solvent system (*MW*_*obs*_ = 113,066 *Da*, Figure 2D). Overall, the charge state distributions observed in the native MS spectra suggest that a substantial portion of HalM2 indeed survives the ionization process in a folded conformation – opening the possibility of studying functionally relevant structural properties of this lanthipeptide synthetase in the gas phase.

To probe the putative native fold of HalM2 in more detail, we performed travelling wave ion mobility^28–30^ measurements on the 20^+^ − 23^+^ ion series (Figure 2E). In these experiments, the HalM2 protein ions are maintained at low kinetic energy while a travelling electromagnetic wave is used to pass the ions through a drift cell filled with nitrogen bath gas. Low energy collisions between the protein and the bath gas molecules impede the transit of the protein ions through the drift cell in a manner that depends on the charge and shape (conformation) of the ion. Ions that are more compact move through the mobility cell more quickly, whereas ions that are more extended move through the cell more slowly. As such, ion mobility (IM) separations allow the conformational landscape of a folded protein to be investigated. Interestingly, the ion mobility drift time distributions of the natively folded HalM2 ions show the presence of two major conformations (Figure 2E). The distributions were fitted with a sum of Gaussian-shaped peaks as described in the Supporting Information to estimate the drift times and fractional abundance of each conformation (Table S3). The drift times for each conformation decrease at higher charge states because more highly charged ions are accelerated to a greater extent by the weak bias voltage (3 V) maintained across the IM cell. Moreover, the relative abundance of the open conformation increases at higher charge states, likely due to electrostatic repulsion of charged amino acid groups on the protein surface. The drift times extracted from the ion mobility data were then used to calculate rotationally averaged collisional cross sections (Ω) for each conformation as described previously^31^ using a set of protein standards of known nitrogen Ω_*N2*_ values^32^ (see Supporting Information). The collisional cross sections for the closed and open conformations (Ω_*c*_ = 63.1 ± 0.7 *nm*^2^ and Ω_*0*_ = 66.9 ± 0.8 *nm*^2^, respectively, Table S4) are in close agreement with the cross section calculated from the HalM2 homology model^21^ using the projection approximation method of Bleiholder (64.2 *nm*^2^).^33^ Thus, HalM2 appears to maintain an ensemble of tertiary structures in the gas phase that closely resembles the overall shape of the molecule predicted by the HalM2 homology model.^21^ This result strongly suggests that the enzyme indeed maintains a near native fold in the gas phase. Similar native IM-MS studies performed on a small collection of other LanM enzymes also provided evidence for the existence of multiple folded gas phase conformations for each of these enzymes (Figure S4). Thus, the gas phase conformational distribution observed for HalM2 may reflect general structural properties of the class II lanthipeptide synthetase fold. Finally, control experiments indicated that the HalM2 conformational distribution was stable over a large range of instrumental settings (Figure S5), suggesting that nESI-IM-MS measurements will likely be feasible on many other lanthipeptide synthetases and RiPP biosynthetic enzymes.

### HalM2 maintains native interactions with the HalA2 precursor peptide in the gas phase

If the HalM2 conformations detected by the ion mobility studies represent natively folded conformations that exist in solution, then these conformations should remain competent for binding to the HalA2 precursor peptide. Accordingly, we tested whether the non-covalently bound Michaelis complex between HalM2 and HalA2 could be kept intact in the gas phase. In the presence of HalA2, the HalM2:HalA2 complex (*MW*_*obs*_ = 120,549 ± 20 *Da*, *MW*_*calc*_ = 120,319 *Da*, Table S1) was clearly detectable as a second set of signals interspersed among the m/z signals for natively folded HalM2 (Figure 3A). Interestingly, the difference between the measured molecular weights for the solvated HalM2 and HalM2:HalA2 Michaelis complex under our conditions (Δ*m*⁄*z* = 7262 *Da*) almost exactly matches the calculated monoisotopic mass of HalA2 (7261.48 *Da*), suggesting that the free enzyme and enzyme:peptide complex ions remain solvated to similar extents. This observation suggests that HalA2 binding might not be associated with a large-scale conformational change in HalM2, or that the sites of residual protein ion solvation are distinct from sites that interact with the precursor peptide. Intriguingly, at higher HalA2 concentrations, weak signals could also be detected for a complex with a stoichiometry of HalM2:(HalA2)2 (Figure 3A, red circles). Thus, HalM2 either possess a 2^nd^ low affinity binding site for HalA2, or the nanoESI process results in non-specific complex formation at higher HalA2 concentrations.

**Figure 3.**
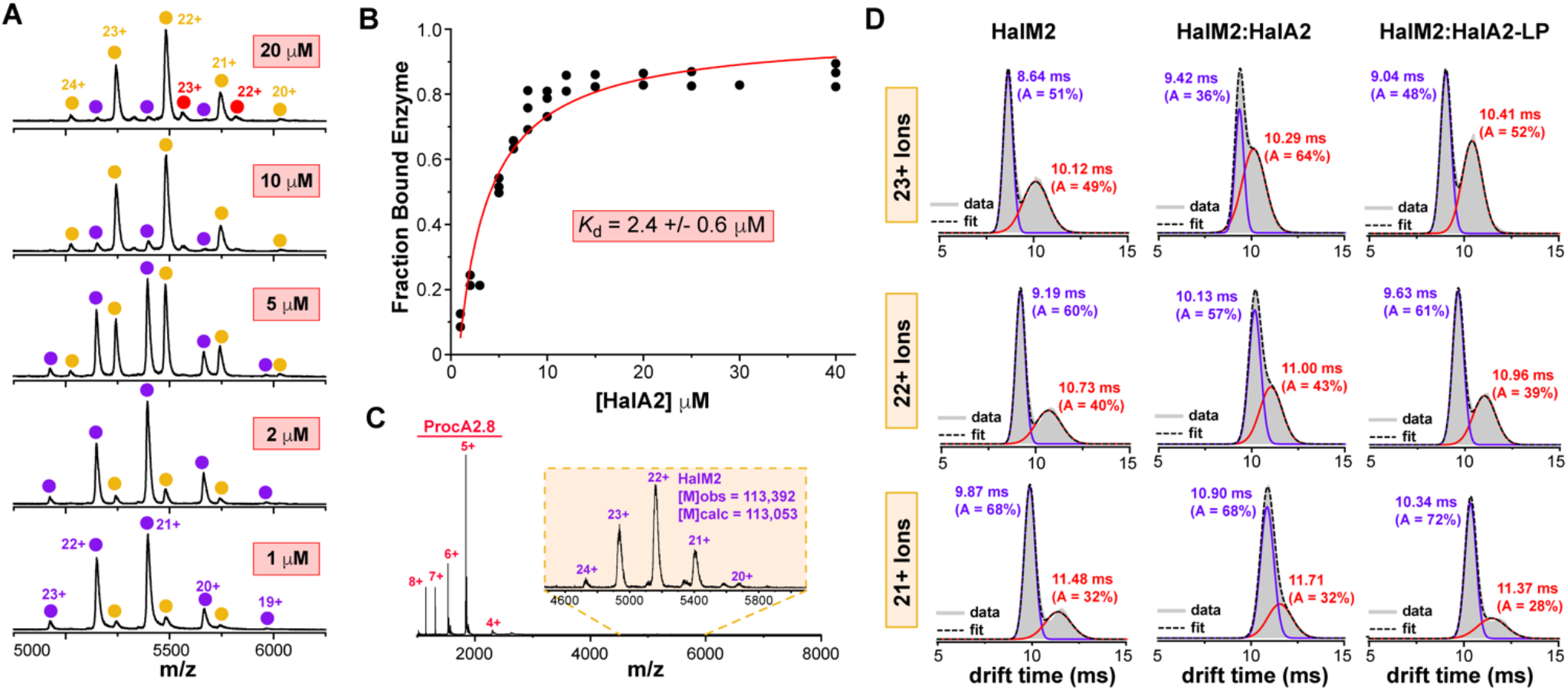
HalM2 maintains native intermolecular contacts with the HalA2 precursor peptide in the gas phase. A) Native nanoESI spectra of HalM2 recorded in the presence of different concentrations of HalA2. As the concentration of HalA2 is increased, the ion signals corresponding to HalM2 (purple dots) shift to a set of signals corresponding to the HalM2:HalA2 complex (gold dots). At higher concentrations, small quantities of a HalM2:(HalA2)2 complex can be detected (red dots). B) Integration of ion peak areas from data such as that shown in panel A allowed a binding affinity for the HalM2:HalA2 complex to be determined (*K*_*d*_ = 2.4 μ*M*). C) HalM2 binding to the non-cognate class II lanthipeptide precursor, ProcA2.8, could not be detected under our conditions, further suggesting that the HalM2:HalA2 complex detected in panel A involves specific molecular recognition mediated by native binding interactions. D) Comparison of ion mobility drift time distributions for HalM2 and its non-covalent complexes with HalA2 and the HalA2 leader peptide (HalA2 residues Met1-Gly36). Distributions were fitted with a sum of two Gaussians to extract the drift times and relative peak areas (Table S3). These drift times were used to calculate nitrogen collisional cross sections for each conformation (Ω_*c*_ and Ω_*0*_ Table S4). Native mass spectra for the HalM2:HalA2 leader peptide complex are shown in Figure S6.

To probe the binding specificity of the complex in more detail, we performed nanoESI experiments in the presence of 20 μM ProcA2.8 (a precursor lanthipeptide from an unrelated class II lanthipeptide biosynthetic pathway).^34^ These experiments failed to provide intense native MS signals for a non-covalent complex between ProcA2.8 and HalM2 (Figure 3C), suggesting that the observed 1:1 complex between HalM2 and HalA2 is not an artifact of the ionization process, but rather involves specific molecular recognition between the synthetase and its cognate precursor peptide. In support of this hypothesis, the MS signals for the HalM2:HalA2 complex could be saturated as a function of HalA2 concentration with a binding affinity (*K*_*d*_ = 2.4 ± 0.6 μ1*M*, Figure 3B) that closely matches the value measured in solution (0.8 μM).^21^ The slight discrepancy between these *K*_*d*_ values may indicate that HalM2:HalA2 binding is driven largely by hydrophobic interactions, which are known to weaken in the gas phase.^35–36^ Consistent with this claim, a recent STD-NMR study of the HalM2-HalA2 interaction suggested that hydrophobic residues in the HalA2 leader peptide are critical for binding to HalM2.^37^ A binding mode involving mainly hydrophobic interactions would also be consistent with the similar solvation levels of the free enzyme and HalM2:HalA2 complex observed in our studies.

### HalA2 binding alters the conformational landscape of HalM2

Ion mobility experiments performed on the HalM2:HalA2 complex again revealed a bimodal drift time distribution suggestive of two main conformational populations with collisional cross section values of Ω_*c*_ = 66.4 ± 0.7 *nm*^2^ and Ω_*0*_ = 68.7 ± 0.9 *nm*^2^ for the compact and open conformations, respectively (Figure 3D, Table S4). The collisional cross section difference between the open and closed conformations of the HalM2:HalA2 complex (ΔΩ = 2.3 *nm*^2^) is less than the difference measured for the free enzyme (ΔΩ = 3.8 *nm*^2^). Thus, peptide binding shifts the HalM2 conformational equilibrium towards a more homogeneous population of structures, perhaps involving the ordering of certain HalM2 structural elements. This effect of peptide binding on the conformational landscape of HalM2 is consistent with the previously proposed conformational selection model for LanM enzymes,^38–41^ wherein binding of the HalA2 leader peptide to an active form of the synthetase shifts the equilibrium population of enzyme conformers towards a more active state. Interestingly, when HalM2 was incubated with the isolated HalA2 leader peptide (HalA2 residues Met1-Gly36), the collisional cross section difference between the open and closed conformations (ΔΩ = 3.3 *nm*^2^) more closely resembles the value measured for the free enzyme (Figure 3D, Table S4). Thus, in the context of the full-length HalA2 precursor, the HalA2 *core peptide* (which contains the sites of post-translational modification) must somehow be directly influencing the tertiary structure of HalM2. These data suggest that nanoESI-IM-MS may prove to be a very useful tool for interrogating RiPP core peptide-synthetase interactions.

To assess the effects of peptide binding on the stability of the gas phase HalM2 ions, we turned to collision induced unfolding (CIU, Figure 4).^42^ This experiment is performed by accelerating the folded protein ions into an argon-filled trap located immediately upstream of the ion mobility mass analyzer. Collisions with the argon molecules thermally excite the folded protein ions into vibrationally excited states that trigger unfolding. The degree of unfolding is then assessed by the ion mobility mass analyzer located immediately downstream of the collision cell in the ion path. These CIU experiments revealed a similar unfolding pattern for HalM2 and the HalM2:HalA2 complex, where the native conformations present under gentle activation conditions (trap collision energy < 40 V) unfold through a series of intermediates (40 V < trap collision energy < 70 V) to more highly denatured conformations (trap collision energy > 70 V). The CIU_50_ values determined from this data (which approximate the collision energy at which half of the protein is unfolded)^42–44^ indicate a substantial stabilization of the HalM2 gas phase structure induced by HalA2 binding 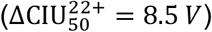. Therefore, the non-covalent contacts between the peptide and synthetase appear to be extensive and are perhaps localized in labile regions of the enzyme that are more susceptible to unfolding in the gas phase. We envision that similar CIU experiments could be very useful for characterizing the binding interfaces between RiPP biosynthetic enzymes and their precursor peptides, and for investigating how post-translational modifications in the peptide may alter enzyme conformation and stability.

**Figure 4.**
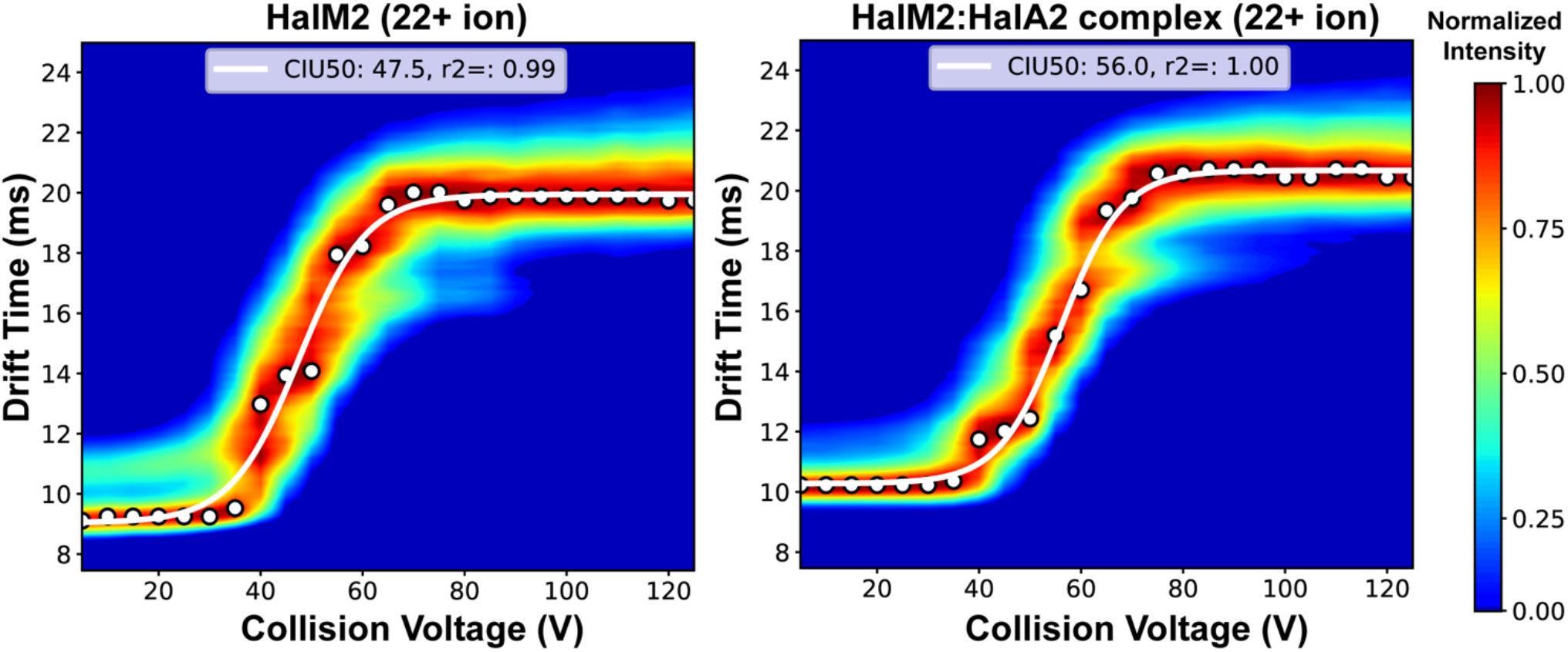
Collision induced unfolding (CIU) analysis of HalM2 and the HalM2:HalA2 Michaelis complex. Unfolding of the 22^+^ ions of HalM2 (left) and its complex with HalA2 (right) as a function of trap collision energy revealed a stabilization of the gas phase enzyme structure induced by HalA2 binding. The unfolding transitions were approximated with CIU_50_ values of 47.5 V and 56.0 V for HalM2 and HalM2:HalA2, respectively, using CIUSuite2 software as described in the Supporting Information.^44^ CIU data were collected with the instrumental settings given in Table S6.

### The HalA2 precursor peptide also possess tertiary structure in the gas phase

Interestingly, during the course of the native mass spectrometry studies of the HalM2:HalA2 complex, we noticed that the HalA2 precursor peptide also exhibited tertiary structure in the gas phase (Figure 5). The mass spectrum of 5 μM HalA2 nanoelectrosprayed from 200 mM ammonium acetate is shown in Figure 5A, along with IM drift time profiles of the 4^+^ to 7^+^ HalA2 ions as a function of trap collision energy (Figure 5B). The data reveal a rich collection of HalA2 conformations that exhibit the expected trend as a function of trap collision energy: namely, the more compact peptide conformations (lower IM drift times) are converted into more open conformations (longer IM drift times) at higher collision energies. As we observed with HalM2, control experiments demonstrated that the HalA2 peptide conformational landscapes were stable over a range of instrumental conditions (Figure S7). The 4^+^ ion exhibited only a single conformation across the entire trap collision energy range tested (up to 100 V), while the more highly charged ions of HalA2 exhibited multiple conformations. Similar IM studies on the HalA2 leader peptide (HalA2 residues Met1-Gly36) demonstrated that at least some of the gas phase structure observed for the full-length HalA2 peptide is attributable to intramolecular interactions within the HalA2 leader peptide (Figure S8). The tertiary structure observed for HalA2 by ion mobility could reflect intramolecular contacts (e.g. salt bridges between charged amino acid side chains) that form once the ions are in the gas phase rather than peptide conformations that exist in solution prior to electrospray. To distinguish among these two possibilities, we performed IM measurements on HalA2 nanoelectrosprayed from a denaturing solvent composed of 50% acetonitrile containing 0.1% formic acid (pH 3.5). This experiment resulted in similar IM drift time distributions for the HalA2 4^+^ to 7^+^ ion series (Figure 5), suggesting that the HalA2 tertiary structure observed in our experiments could be reporting on gas phase conformational changes, rather than on higher order structure that pre-existed in solution. Thus, while these data ultimately failed to provide conclusive evidence for the existence of a structured HalA2 precursor peptide in solution, the data nevertheless reveal the potential of native nanoESI-IM-MS for investigating the presence of higher order structure that may exist in other RiPP precursor peptides. The sequence diversity of lanthipeptide precursors is vast,^45^ and aside from a handful of literature reports,^46–49^ the structural properties of lanthipeptide and other RiPP precursors have not been extensively studied.

**Figure 5.**
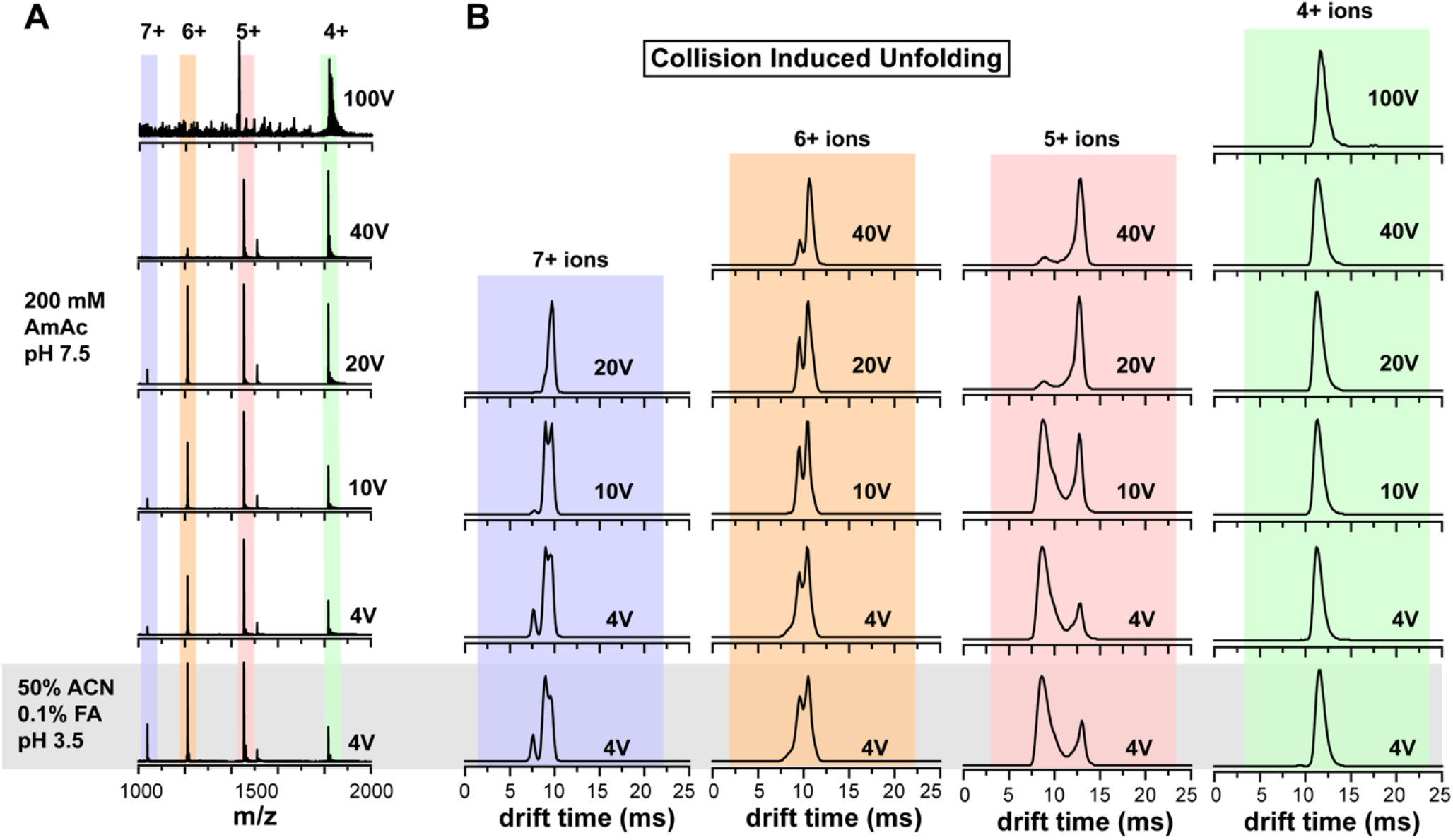
Native nESI-IM-MS analysis of the HalA2 precursor peptide reveals a rich conformational distribution. A) Mass spectra of 5 μM HalA2 nanoelectrosprayed from either 200 mM ammonium acetate (pH 7.5) or 50% acetonitrile, 0.1% formic acid (pH 3.5). In addition, the mixtures were spiked with 100 μM TCEP to ensure complete reduction of the cysteine residues in the peptide. The HalA2 mass determined from the native mass spectrum, 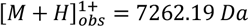, is in close agreement with the calculated monoisotopic mass of the fully reduced peptide, 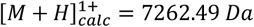. B) Ion mobility drift time distributions for the indicated HalA2 ions as a function of trap collision energy. Data were collected using the instrumental settings given in Table S7. The trap collision energy used to acquire the data is indicated in each panel. More highly charged HalA2 ions fragment at lower collision energy, precluding the collection of ion mobility data above collision energies of 20 V (for the 7^+^ ions), 40 V (for the 6^+^ and 5^+^ ions), and 100 V (for the 4^+^ ion).

### Mutations in structurally dynamic regions of HalM2 alter the conformational landscape and its sensitivity to HalA2 binding

We have recently reported a hydrogen-deuterium exchange mass spectrometry study of HalM2,^21^ wherein a number of HalM2 structural elements (Figure 6A-6B) were found to exhibit reduced solvent deuterium uptake into the amide moieties of the protein backbone upon binding of the enzyme to nucleotide or HalA2. These data are consistent with a structural organization within these elements of HalM2 upon ligand binding. Replacement of these elements with Gly-Ser linkers resulted in various perturbations to HalM2 substrate binding and catalysis, validating their important roles in the enzymatic function of HalM2.^21^ In the present study, we employed our optimized nESI-IM-MS conditions to evaluate the tertiary structures and conformational distributions of these variant enzymes in the presence and absence of the HalA2 precursor peptide. Native nESI mass spectra recorded in the presence of HalA2 are shown for each enzyme in Figure 6C, along with ion mobility drift time distributions extracted for the 22^+^ charge states of the free enzymes (Figure 6D) and enzyme:peptide complexes (Figure 6E). As observed for wild type (wt) HalM2, the ion mobility data for each HalM2 variant provided evidence for two major gas phase populations that could be well-fitted with a sum of two Gaussian-shaped peaks (Table S3).

**Figure 6.**
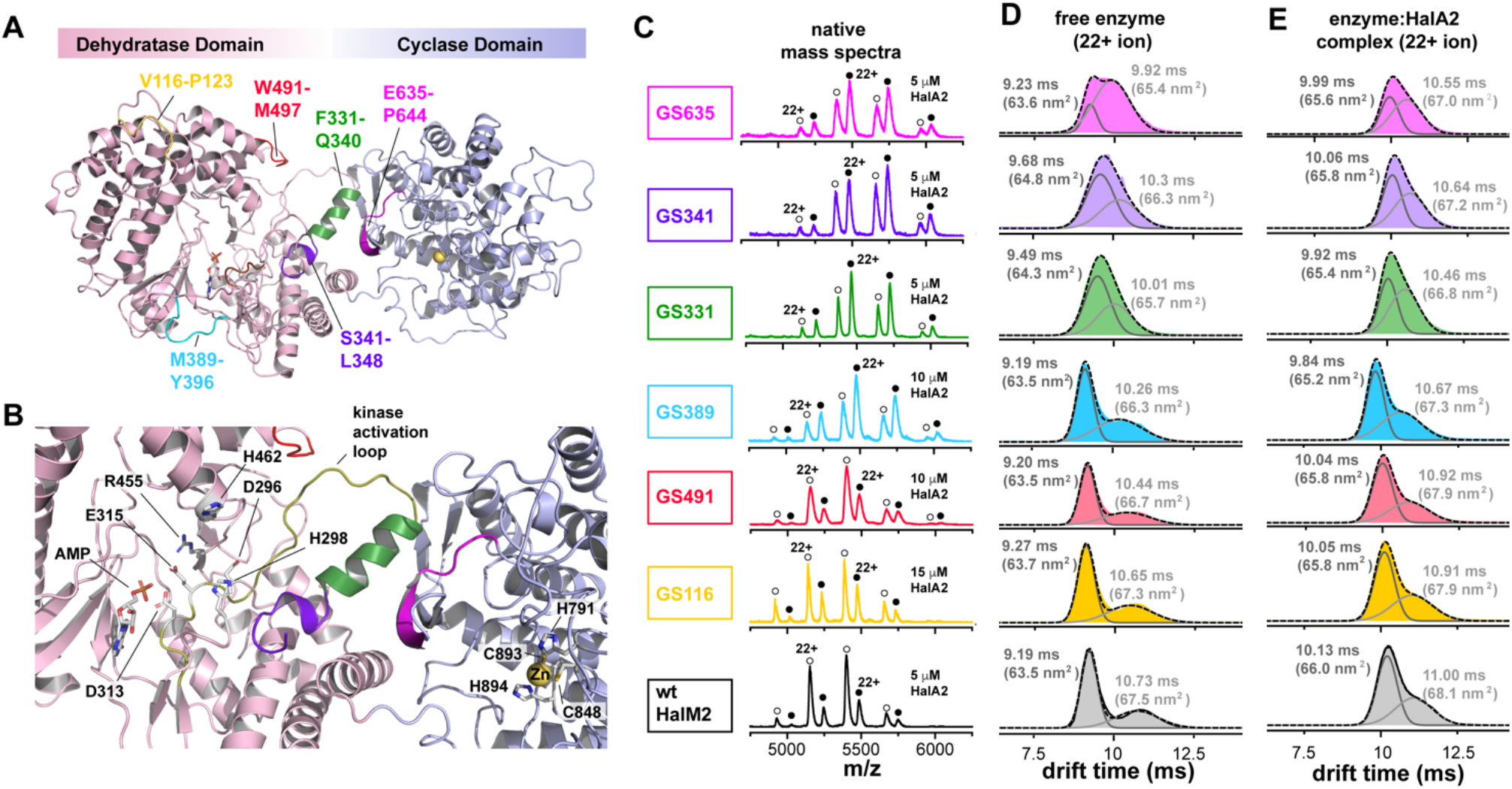
Native nESI-IM-MS characterization of HalM2 variant enzymes. A) HalM2 homology model built from the X-ray crystal structure of CylM.^9^ Dynamic HalM2 structural elements that undergo an organization (i.e. less deuterium uptake in HDX-MS studies) upon HalA2 binding are indicated.^21^ These segments of HalM2 were replaced with Gly-Ser linkers as described previously.^21^ B) Closer view of critical structural elements at the interface of the dehydratase C-lobe and the cyclase domain: F331-Q340 (green), S341-L348 (violet), and E635-P644 (fuchsia). The unstructured P349-P405 loop has been removed for clarity. Residues that are essential for dehydration (D296, H298, D313, E315, R455, H462) and cyclization (H791, C848, C893, H894) catalysis are shown, as are the AMP molecule and Zn^2+^ ion bound in the dehydratase and cyclase active sites. C) Native mass spectra of wt HalM2 and its variant enzymes bound to HalA2. The charge state distributions corresponding to the free enzymes and enzyme:HalA2 complexes are denoted with the open and filled circles, respectively. The 22^+^ ions used to compare the drift time profiles and the HalA2 concentrations used to generate the spectra are indicated in each plot. Ion mobility drift time distributions are shown in panels (D) and (E) for the 22^+^ ions of the free enzymes (panel D) and enzyme:HalA2 complexes (panel E). The distributions have been fitted with a sum of two Gaussians to extract the drift times and collisional cross sections of the two conformations. A summary of the Gaussian fits and CCS calculations for each charge state of all enzymes and their HalA2 complexes are provided in Tables S3 and S4, respectively.

The drift time profiles of the GS116, GS491, and GS389 enzymes revealed conformational distributions both in the presence and absence of HalA2 that closely matched the wt HalM2 enzyme. Thus, any substrate binding or catalytic defects induced by these mutations likely originate from local structural perturbations that do not greatly impact the global structure of the enzyme. The GS116 mutation (replacement of V116-P123 with a (GS)4 linker) in the helical capping domain of the dehydratase (Figure 6A) has been shown to weaken the HalA2 binding affinity by approximately 16-fold, without otherwise strongly affecting enzymatic catalysis.^21^ The ion mobility data for GS116 suggests that this mutation likely induces a local structural perturbation that is detrimental to HalA2 binding, rather than a perturbation on the overall HalM2 fold. The GS389 mutation (replacement of M389-Y396 with a (GS)4 linker) involves removal of a putative β-strand within a large mobile loop (HalM2 residues P349-P405) that is not ordered in the crystal structure of the LanM homologue used to generate the HalM2 homology model.^9^ The GS389 variant has a 6-fold weakened HalA2 binding affinity and a significant defect in the phosphate elimination step of the dehydration reaction (see Figure 1B). These data led us to propose that the M389-Y396 region may assist in forming a HalM2 conformation that facilitates peptide docking into the dehydratase active site and promotes phosphate elimination. The ion mobility data suggest that any structure formed by the putative M389-Y396 *β*-strand must either be highly localized, transiently formed during turnover, and/or unstable in the gas phase so as not to greatly influence the overall shape of the enzyme. Finally, the GS491 mutation (replacement of W491-M497 with a (GS)4 linker) is located at the interface of the kinase activation domain and the kinase activation loop (Figure 6A). This mutation leads to a 5-fold reduction in HalA2 binding affinity and was shown reduce the rates of both HalA2 dehydration and cyclization.^21^ Again, the ion mobility data suggest that these catalytic defects appear to arise from local and/or transient structural perturbations that do not greatly alter the conformational landscape of the enzyme.

In contrast to the variants discussed above, the GS331, GS341, and GS635 mutations (replacement of HalM2 residues F331-Q340, S341-L348, and E635-P644 with (GS)_5_, (GS)_4_, and (GS)_5_ linkers, respectively, Figure 6A-6B) each perturb the overall conformational landscape of HalM2. Namely, in the free enzymes, the collisional cross section difference between the open and compact conformations ranges from 1.4 to 1.8 nm^2^ (ΔΩ_*free*_, Table 1). In contrast, the ΔΩ_*free*_ values for the other variants range from 2.8 to 4.0 nm^2^ and lead to the two readily distinguishable conformational populations of the free enzyme. Second, upon binding to HalA2, the ΔΩ_*complex*_ values for the GS331, GS341, and GS635 variants are either unaltered or are not greatly affected relative to ΔΩ_*free*_ (Table 1), suggesting that HalA2 binding is not triggering the same level of structural organization that is observed in the other enzymes. The structural elements mutated in GS331, GS341, and GS635 comprise helical regions in the C-lobe of the dehydratase (F331-Q340 and S341-L348) and a *β*-stranded region in the cyclase domain (E635-P644) that appear to closely interact in the HalM2 homology model (Figure 6B). Apparently, mutation of any one of these elements affects the enzyme conformational landscape and its ability to respond to HalA2 binding. Our previous HDX-MS studies suggested that while the F331-Q340, S341-L348, and E635-P644 elements each become more structured upon HalA2 binding,^21^ none of these mutations significantly alters HalA2 binding affinity, and each mutation exerted significant effects on HalM2 catalysis. Thus, HalA2 binding could potentially alter the structure and dynamics of these elements through allosteric effects, which may be important for ensuring high-level enzyme activity. Cumulatively, the data provide strong support that the integrity of the interface between the dehydratase and cyclase domains (as mediated by F331-Q340, S341-L348, and E635-P644) strongly influences the conformational landscape of the enzyme and is critical for the structure and function of HalM2.

**Table 1.**
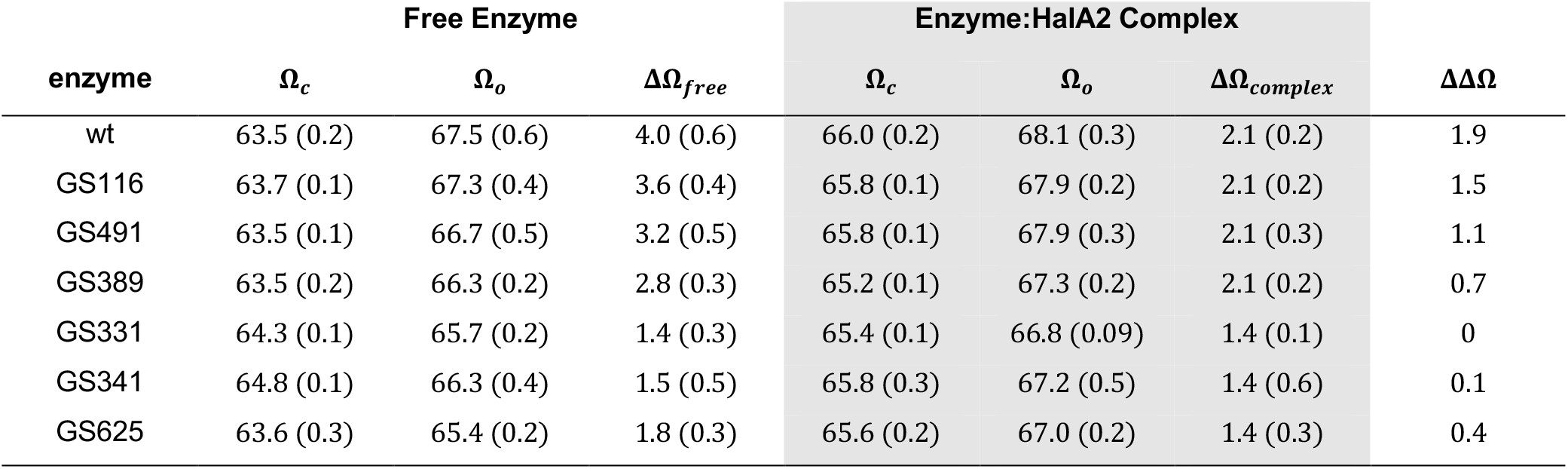
Collisional cross sections (in units of *nm*^2^) for the 22^+^ ions of the compact (Ω_*c*_) and open (Ω_*0*_) conformations of the enzyme and enzyme:HalA2 complexes. Ion mobiligrams were acquired in triplicate and were fitted with a sum of two Gaussian-shaped peaks to extract the drift times for the two conformations, which were then used to calculate the Ω_*c*_ and Ω_*0*_ values (Tables S3 and S4). The standard error of the mean is reported in parenthesis. Ω_*c*_ and Ω_*0*_ were then used to calculate ΔΩ_*free*_, ΔΩ_*complex*_, and ΔΔΩ = ΔΩ_*free*_ − ΔΩ_*complex*_.

| enzyme | Free Enzyme | | | Enzyme:HalA2 Complex | | | $\Delta\Delta\Omega$ |
| --- | --- | --- | --- | --- | --- | --- | --- |
| | $\Omega_c$ | $\Omega_o$ | $\Delta\Omega_{free}$ | $\Omega_c$ | $\Omega_o$ | $\Delta\Omega_{complex}$ | |
| wt | 63.5 (0.2) | 67.5 (0.6) | 4.0 (0.6) | 66.0 (0.2) | 68.1 (0.3) | 2.1 (0.2) | 1.9 |
| GS116 | 63.7 (0.1) | 67.3 (0.4) | 3.6 (0.4) | 65.8 (0.1) | 67.9 (0.2) | 2.1 (0.2) | 1.5 |
| GS491 | 63.5 (0.1) | 66.7 (0.5) | 3.2 (0.5) | 65.8 (0.1) | 67.9 (0.3) | 2.1 (0.3) | 1.1 |
| GS389 | 63.5 (0.2) | 66.3 (0.2) | 2.8 (0.3) | 65.2 (0.1) | 67.3 (0.2) | 2.1 (0.2) | 0.7 |
| GS331 | 64.3 (0.1) | 65.7 (0.2) | 1.4 (0.3) | 65.4 (0.1) | 66.8 (0.09) | 1.4 (0.1) | 0 |
| GS341 | 64.8 (0.1) | 66.3 (0.4) | 1.5 (0.5) | 65.8 (0.3) | 67.2 (0.5) | 1.4 (0.6) | 0.1 |
| GS625 | 63.6 (0.3) | 65.4 (0.2) | 1.8 (0.3) | 65.6 (0.2) | 67.0 (0.2) | 1.4 (0.3) | 0.4 |

## Conclusion

In this work, we have provided to our knowledge the first detailed characterization of a RiPP biosynthetic enzyme using native nanoelectrospray ionization coupled to ion mobility mass spectrometry (nanoESI-IM-MS). Through a series of careful validation and control experiments, we have shown that the class II lanthipeptide synthetase, HalM2, maintains a near-native fold in the gas phase of a commercially available mass spectrometer over a large working range of instrumental parameters. This assertion is strongly supported by the close agreement between the collisional cross sections calculated from the ion mobility data and the HalM2 homology model, and by the specificity, stoichiometry, and binding affinity determined for the gas phase HalM2:HalA2 complex. The nESI-IM-MS approach has also revealed a number of intriguing observations that provide further support for existing models of LanM function. For example, the bimodal drift time distribution observed for HalM2 and the sensitivity of this distribution to HalA2 binding (Figure 3D) is consistent with the previously proposed conformational selection model for LanM function.^38^ In addition, the differential responses of the HalM2 conformational landscape to full length HalA2 and to the isolated HalA2 leader peptide (Figure 3D), provide support for a direct interaction between the synthetase and the core peptide in a manner that influences the topology of the synthetase. Interestingly, mutations to conserved structural elements near the interface of the dehydratase and cyclase domains also perturb the conformational landscape of the free enzyme and diminish the effects of peptide binding on the collisional cross section difference of the two major enzyme conformations (the ΔΔΩ parameter, Table 1). Thus, the mutated elements (the F331-Q340 and S341-L348 helices of the dehydratse domain and the E635-P644 *β* strand of the cyclase domain, Figure 6) appear to play an essential role in defining the conformational landscape and in transmitting structural perturbations induced by HalA2 binding that are essential for efficient catalysis.^21^ The similar residual solvation of the free HalM2 and HalM2:HalA2 ions suggests that HalA2 binding is not likely to be accompanied by a large conformational change in the enzyme. However, binding of HalA2 nevertheless substantially stabilizes HalM2 towards collision induced unfolding (Figure 4), suggesting that the peptide may bind to an extended interface across the enzyme surface. It is tempting to speculate that this stabilization may derive in part from interactions between the HalA2 peptide with the interface of the dehydratase and cyclase domains, where the F331-Q340 helix and E635-P644 *β*-strand interact (Figure 6B). Consistent with this hypothesis, recent photochemical crosslinking studies have provided evidence for direct interactions between the HalA2 leader peptide with the cyclase domain in the vicinity of the E635-P644 *β*-strand, as well as with the *N*-terminal capping domain in the vicinity of V116-P123.^37^ Given the apparent spatial distance separating these two elements in the HalM2 homology model (Figure 6A), it is possible that HalM2 possess two separate binding sites for HalA2. Curiously, we were able to detect small quantities of a HalM2:(HalA2)_2_ complex (Figure 3A) and a HalM2:(HalA2-LP)2 complex (Figure S6) in our titration studies at higher peptide concentrations. Altogether, the present study illustrates the potentially broad utility of native nESI-IM-MS for studying conformational changes and non-covalent enzyme:peptide interactions in RiPP biosynthetic systems. Combined with our previous HDX-MS analysis on HalM2,^21^ it is evident that mass spectrometry-based structural biology approaches fill an important niche in the mechanistic characterization of RiPP biosynthesis and allow questions regarding the relevance of conformational dynamics in these systems to be addressed.

## Methods

### Preparation of HalM2, its variant enzymes, and HalA2 for native MS

Wild type (wt) HalM2 and its variant enzymes were cloned, expressed, and purified as described previously.^21^ The night before native MS analysis, aliquots of the enzyme were removed from the freezer and were treated with thrombin (0.1 kU/mL, Promega) overnight at 4 °C in enzyme storage buffer (20 mM HEPES, 300 mM KCl, 10 % glycerol, pH 7.5) to remove the *N*-terminal His_6_ tag used for affinity purification. Immediately prior to native MS analysis, the cleaved His_6_ tag was removed and the enzymes were buffer exchanged into 200 mM ammonium acetate (pH 7.5) using Micro Bio-Spin 6 columns (Bio-Rad). His_6_-affinity tagged HalA2 was likewise digested with thrombin to remove the His_6_ tag, purified by reverse phase HPLC (see supporting methods), lyophilized to dryness, and resuspended in 200 mM ammonium acetate for native MS studies. Immediately prior to use, the concentrations of enzyme and HalA2 samples were determined from their absorption at 280 nm using extinction coefficients calculated from the enzyme/peptide sequence using the ExPASy ProtParam tool and were then diluted to the desired values in 200 mM ammonium acetate (pH 7.5).

### Preparation of platinum coated borosilicate nanospray emitters

The nanospray emitters used for all native mass spectrometry experiments were made in house by pulling borosilicate glass capillaries (1.0 OD × 0.78 ID × 100 L mm, Harvard Apparatus GC100T-10) with a Sutter Instrument Model P-2000 tip puller using the following instrumental parameters: Heat = 350, Filament = 4, Velocity = 60, Delay = 255, Pull = 0. The diameter of the tips prepared in this way were estimated to be approximately 1-2 μM by light microscopy. The tips were then coated with 10 nm platinum at a height of 7 mm using a Leica Microsystems EM ACE600 High Resolution Sputter Coater at the Facility for Electron Microscopy Research at McGill University.

### Mass spectrometry

Typically, 5-10 μL of sample for native MS was loaded into a nanospray emitter using Eppendorf 20 μL GELoader tips. Loaded tips were fastened into a tip holder (Thermo Scientific part ES286), briefly centrifuged for 2-3 s to move the fluid to the end of the tip and were mounted onto a Synapt G2-Si ion mobility mass spectrometer (Waters) equipped with a nanospray ESI source. All MS data were acquired in positive ion and sensitivity modes over an m/z range from 100-8000 using a 1 s scan rate. The TOF detector was calibrated over the same m/z range with sodium iodide. Unless otherwise noted, the standard “gentle” ionization conditions were employed (Table S5). Key setting include the capillary voltage = 1.5 kV, cone voltage = 1 V, source offset = 1 V, source temperature = 35 °C, trap collision energy = 4 V, trap gas (argon) flow = 2 mL/min, and trap DC bias = 35 V. For ion mobility measurements: IMS gas (nitrogen) flow = 90 mL/min, IMS bias = 3 V, IMS wave velocity = 550 m/s, and wave height = 40V. For the collision induced unfolding studies (Figure 4), the cone voltage and source offset were increased to 50 V, the source temperature was increased to 70 °C, the trap gas flow was increased to 10 mL/min, and the trap DC bias was increased to 45 V while the trap collision energy was varied from 5-180 V (Table S6). These altered instrumental settings were necessary for maintaining sufficient ion transmission at the higher trap CE settings used in the CIU experiemtn. For ion mobility studies of HalA2 and the HalA2 leader peptide (Table S7), the only changes to the standard settings were the source temperature (70 °C), trap DC bias (30 V), IMS wave velocity (400 m/s) and IMS wave height (25 V). All data were processed using MassLynx (Waters), Microsoft Excel, and OriginPro as needed.

## Supporting Information Available

Purification protocols for enzymes and peptides, calibration and data fitting procedures, additional mass spectra, descriptions of biochemical assays, tables of calculated collisional cross section values, and complete lists of instrumental conditions are available free of charge.

## Acknowledgement

The authors thank Ms. Annie Shao and Ms. Christine Traaseth for assisting with the preparation of some of the enzymes. This work was funded by a Natural Sciences and Engineering Research Council of Canada Discovery Grant (RGPIN/04485-2017), the Canada Foundation for Innovation, the Fonds de recherche du Québec – Santé, and the Government of Canada’s New Frontiers in Research Fund. YH was supported by a fellowship from the Centre du Recherche en Biologie Structurale (CRBS).

---

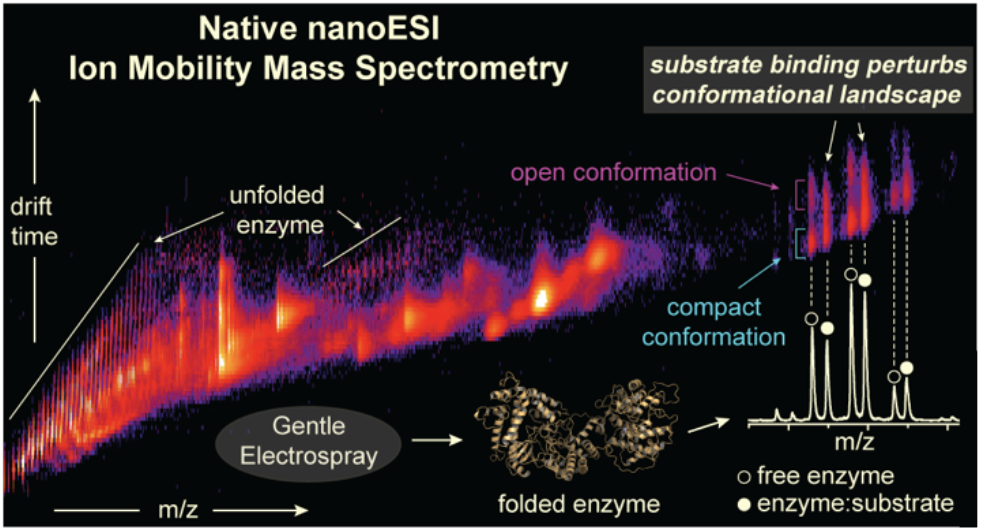
For Table of Contents Only.

