## Supporting Information for "Exploring the conformational landscape of a lanthipeptide synthetase using native mass spectrometry"

### Supporting Methods

_Expression of $His_6$-HalA2, $His_6$-HalM2 and HalM2 variants:_ The *N*-terminally $His_6$-tagged HalA2 peptide sequence was cloned into the *pET15* vector and was transformed into *E. coli* BL21 (DE3) cells. Transformed cells were plated on LB-agar containing ampicillin (100 µg/mL). A single colony was cultured in 5 mL of LB supplemented with 100 µg/mL ampicillin at 37 °C with shaking overnight. A 600 µL portion of this overnight culture was used to grow a second culture of 60 mL as described above. After 12-15 hours, the culture was centrifuged at 4700 rpm for 10 minutes and the supernatant was discarded in order to remove any β-lactamase that was produced. The pellet was re-suspended in a 60 mL of fresh LB. This was then used to inoculate 6 L of fresh LB supplemented with 100 µg/mL ampicillin. The inoculated LB was grown at 37 °C with shaking at 160 rpm until the $OD_{600}$ reached 0.75-0.85. IPTG was added to a final concentration of 1 mM and the bacteria were grown for another 3 hours at 37 °C. When expressing HalM2 and its variants, after IPTG induction, bacteria was grown overnight at 18 °C. Cells were harvested by centrifugation (5000 X *g* for 20 minutes at 4 °C) and the cell pellets were stored at -80 °C until purification.

_HalA2 purification:_ The frozen cell pellet was suspended in wash buffer (6 M guanidine hydrochloride, 20 mM $NaH_2PO_4$, 500 mM NaCl, 0.5 mM Imidazole pH 7.5) and was sonicated (35% amplitude, 4.0 s pulse on, 9.9 s pulse off) for 5-10 minutes keeping the sample at 4 °C. The lysed cells were then centrifuged at 15000 x *g* for 45 minutes at 4 °C. The subsequent steps were carried out at room temperature. With a 1.5 mL/min flow rate, the supernatant was loaded onto a 5 mL HisTrap FF Ni-NTA affinity column (GE Healthcare) that was charged with 3 column volumes (CV) of 0.1 M $NiSO_4$ and equilibrated with 6 CV of wash buffer. After loading the sample, the column was washed with 6 CV of wash buffer. The peptide was then eluted with a 5 CV of elution buffer (4 M guanidine hydrochloride, 20 mM Tris, 100 mM NaCl, 1 M Imidazole, pH 7.5). Following the Ni-NTA affinity purification, the peptide was desalted using a C4-Solid Phase Extraction column equilibrated with 0.1% trifluoroacetic acid (TFA) with 80% ACN, 0.1% TFA. Purified peptide was then lyophilized before re-dissolving it in 5 mM phosphate buffer (pH 7.5) and 2 mM TCEP. High purity thrombin (MP Biomedicals) was added to a final concentration of 0.1 kU/mL and was kept in a shaker at room temperature overnight in order to remove the *N*-terminal his tag prior to HPLC purification. The sample was centrifuged at 16000 x *g* to remove any precipitate and the supernatant was injected to a C-8 Supelco Reverse Phase Semi-prep column (Sigma Aldrich). Solvent A and B were 0.1 % TFA and 80% ACN with 0.1 % TFA respectively. The peptide was eluted in a solvent gradient from 3 – 100 % B over 35 minutes. Tagless HalA2 eluted at approximately 17 minutes. The purified peptide was lyophilized and re-dissolved in water and stored at -80 °C.

*Activity assays in HEPES and Ammonium acetate buffer*: HalM2 activity was measured either in 100 mM HEPES (pH 7.5) or 200 mM ammonium acetate (pH 7.5) as previously described.[1] Briefly, assays contained 1 μM HalM2, 50 μM HalA2, 1 mM TCEP, 5 mM $MgCl_2$ and 5 mM ATP and were quenched after 8 min by 10 fold dilution into 100 mM citrate (pH 3.5) containing 1 mM EDTA and 10 mM TCEP. Samples were incubated at 25 °C for 10 min, at which point the pH was neutralized to 6.0 by addition of 5 M NaOH. The samples were then spiked with 10 mM N-ethylmaleimide (NEM) to alkylate the free Cys thiols, incubated at 37 °C for 10 min, and acidified by addition of trifluoroacetic acid to 1% (v/v). Quenched samples were desalted using C4-solid phase extraction and lyophilized. For LC-ESI-MS analysis, the lyophilized samples were dissolved in water containing 0.1% formic acid and were separated using a Waters BEH C8 UPLC column and a linear mobile phase gradient of 0-100% solvent B (acetonitrile containing 0.1% formic acid). Solvent A was water containing 0.1% formic acid. The C8 column effluent was directed to the electrospray ionization (ESI) source, where peptides were ionized using a capillary voltage of 3.0 kV, cone voltage of 80 V, source offset of 40 V, and a source temperature of 250 °C. Mass spectral data were collected in positive ion and sensitivity modes over an m/z range from 100-2000 m/z with a 1 s scan rate. An external [Glu1]-fibrinopeptin B standard was used to correct the m/z values. The results from these assays (Figure S1) agree well with previous data on HalM2[1] and suggest nearly indistinguishable activity in the two buffers.

*Determination of drift times from the ion mobility data:* For each enzyme and enzyme:peptide complex, ion mobility-resolved nESI mass spectra were collected in triplicate. From this data, the ion mobility drift time distributions for each charge state of interest were extracted using MassLynx software (Waters). The drift time mobiligrams were then imported into Microsoft Excel, normalized, combined into a single data set, and imported into OriginPro 2018 for fitting. The combined triplicate data sets were then fitted to a sum of Gaussian peaks using forms of Equation 1:

Equation 1:

$$y = y_0 + \sum \frac{A}{w\sqrt{\pi/2}} e^{-2\frac{(x-x_c)^2}{w^2}}$$

Here, $A$ is the area of the peak, $w$ is the peak width, $x_c$ is the drift time, and $y_0$ is a baseline correction. If the $R^2$ value of the initial fit was found to be < 0.99, the data were fitted with additional Gaussian peaks to achieve a better fit. *F*-tests were then used to verify that the parameters introduced by the additional Gaussian term indeed significantly improved the fit (at the $\alpha = 0.05$ level) by reducing the residual sum squared error (data not shown). A summary of all Gaussian fits performed for this study is provided in Table S3.

*Calculation of collisional cross sections from ion mobility drift times*: The drift times reported in Table S3 were then used to calculate collisional cross sections for each enzyme and non-covalent enzyme:peptide complex of interest. The mobility of an ion ($K$) is related to its collisional cross section ($\Omega$) by the Mason-Schamp equation (Equation 2):[2-3]

Equation 2:

$$\Omega = \frac{3ez}{16N}\left(\frac{2\pi}{\mu k_B T}\right)^{1/2}\frac{1}{K}$$

Here, $N$ is the drift gas number density, $e$ is the elementary charge, $z$ is the charge state of the ion, $\mu$ is the reduced mass of the ion of interest and drift gas, $T$ is the drift gas temperature and $k_B$ is the Boltzmann constant. In travelling wave ion mobility (TWIM) experiments such as the studies performed here, the relationship between the measured drift time of an ion and its mobility ($K$) is non-linear and is strongly dependent on the instrumental conditions. Thus, in TWIM experiments, the Ω values of the ions of interest are determined by calibration against a set of standard proteins with known Ω values. For our study, we used concanavalin A (Sigma, C2010), avidin (Sigma, A9275), and alcohol dehydrogenase (Sigma, A7011) as the protein standards. Bush et. al. previously measured nitrogen collisional cross section values ($\Omega_{N2}$) for the various charge states of these standard proteins in 200 mM ammonium acetate using a modified drift tube ion mobility instrument (Table S2), which allowed for a direct determination of $\Omega_{N2}$ from the measured ion drift times.[4] Accordingly, we prepared concentrated stock solutions of these proteins by dissolving the lyophilized powders in 200 mM ammonium acetate. These stock solutions were then further buffer exchanged into fresh 200 mM ammonium acetate using Micro Bio-Spin P6 columns (Bio-Rad) and diluted to a final working concentration of 5 μM in 200 mM ammonium acetate. The standard proteins were subjected to native ESI using instrumental conditions that were identical to those used for collecting the TWIM data for HalM2, its variant enzymes, and their non-covalent Michaelis complexes with HalA2 (Table S5). MS spectra for the standards are shown in Figure S2, along with ion mobiligrams of the charge states that were used to construct the calibration curve.

To construct the calibration curve, the experimentally measured drift times ($t_D$) for the standard proteins were first corrected to account for $m/z$ – dependent flight times according to equation 3:

Equation 3:

$$t'_D = t_D - 1.57\sqrt{m/z}$$

The $m/z$ in Equation 3 corresponds to the observed value measured in the experiment for the ion in question (Table S2), and the factor of 1.57 is an instrumental constant.[5]

Likewise, the known $\Omega_{N2}$ values for the standard proteins were corrected for their reduced mass (with $N_2$) and charge according to Equation 4:

Equation 4:

$$\Omega'_{N2} = \frac{\Omega_{N2}\sqrt{\mu}}{z}$$

A plot of $\ln(t'_D)$ vs. $\ln(\Omega'_{N2})$ was then fitted with linear regression (Figure S3A). The slope term from this fit ($X$) compensates for the non-linear effects of the TWIM analyzer, while the intercept term corrects for the temperature, pressure, and electric field parameters (which remain constant as long as the standard proteins and the proteins of interest are collected with exactly the same instrumental settings). The slope term ($X$) from the fit in Figure S3A is then used to correct the $t'_D$values for the protein standards according to equation 5:

Equation 5:

$$\mathrm{t}''_D = {t'_D}^X\left(\frac{z}{\sqrt{\mu}}\right)$$

Finally, the $\mathrm{t}''_D$ parameter is plotted vs. $\Omega_{N2}$ and is again subjected to linear regression to construct the final calibration curve (Figure S3B). The $\mathrm{t}''_D$ values are then determined from the measured drift times of the unknowns (i.e. the drift time values, $x_c$, derived from the Gaussian fits reported in Table S3) using Equations 3 and 5, and are plugged into the linear function derived from the calibration curve in Figure S4B to determine the corresponding Ω values for the ions of interest that are reported in Tables 1, and S4. In addition, for ease of comparison we also report $\Omega_{avg}$ values, which are simply the arithmetic mean of the respective Ω values measured at each individual charge state for a given protein/protein-peptide complex.

<u>*Effects of instrumental parameters on the HalM2 conformational landscape*</u>: Excessive thermal activation derived from collisions between the protein ions and the gasses

present in the mass spectrometer can lead to the unfolding of native protein structures during transit of the ions from the source to the time-of-flight (TOF) detector. Thus, starting from our standard conditions (Table S5), we carefully examined the effects of various instrumental parameters on the distribution of HalM2 between the open and closed conformations. For these studies, we recorded IM data for HalM2 over a range of temperatures (Figure S5A), cone and source offset voltages (Figure S5B), trap collision energies (Figure S5C), and trap bias voltages (Figure S5D). Below a source temperature of 100 °C, and cone and source offset voltages of 100 V, the drift time distribution of mobility-separated HalM2 ions remained constant, suggesting that the gas phase HalM2 tertiary structure is largely resistant to perturbations in these ion source parameters over a range of operational values. We also subjected the natively-folded HalM2 ions to collision induced unfolding (CIU) in the trap region of the IM cell (Figure S5C).[6] This experiment is performed by accelerating the folded protein ions into an argon-filled trap located immediately upstream of the ion mobility mass analyzer. Collisions with the argon molecules thermally excite the folded protein ions into vibrationally excited states that trigger unfolding. The degree of unfolding is then assessed by the ion mobility mass analyzer located immediately downstream of the collision cell in the ion path. The CIU experiment revealed little change in the HalM2 conformational landscape up to a trap collision energy setting of 30 V. Between 30-40 V of trap collision energy, the more open conformation decreased in intensity, while the drift time of the more compact conformation remained constant. Above 40 V, the compact conformation began to unfold through a series of intermediate states characterized by longer drift times. Finally, we adjusted the trap bias voltage, an instrumental parameter that controls the transfer of ions from the argon trap to the ion mobility cell (Figure S5D). Under our conditions, the HalM2 conformational landscape remained constant up to a trap bias voltage of approximately 85 V, above which, the protein began to unfold. Overall, these data suggest that the major conformational isomers of HalM2 are relatively stable species in the gas phase, and do not appear to interconvert on the time scale of the measurement as long as the instrumental parameters are kept below their threshold values. Therefore, the observed conformations likely reflect a conformational distribution that exists in solution prior to electrospray, or to a distribution that equilibrates rapidly once the protein ions have entered the gas phase. Moreover, these relatively permissive instrumental settings suggest that nESI-IM-MS measurements will likely be feasible on many other lanthipeptide synthetases and RiPP biosynthetic enzymes.

*Collision induced unfolding (CIU) experiments:* The natively folded HalM2 and HalM2:HalA2 complex were subjected to CIU experiments to probe the stability of their gas phase structures (Figure 4, Figure S7). CIU is achieved by accelerating the ions into the argon-filled trap region of the TWIM analyzer. The kinetic energy delivered to the ions during this step can be controlled by incrementally changing the "Trap CE" instrumental parameter on the Waters Synapt G2-Si. Measurements were made in duplicate in positive ion, sensitivity, and mobility TOF modes. Mobility-resolved spectra (m/z range = 100-8000) were collected over a 15 s interval. HalM2 and HalA2 were used at concentrations of 5 and 10 μM (respectively, when present) in 200 mM ammonium acetate. The relevant instrumental settings for the CIU studies are provided in Table S6. Of special note, we found that the trap gas flow had to be increased to 10 mL/min because at the 2 mL/min flow rate that was employed in our standard conditions, the baseline of the ion mobiligram was unacceptably high at trap CE values > 50 V. The trap CE was varied from 5 – 180 V in increments of 5 V to trigger the CIU. Raw drift time data for the $22^{+}$ ions of the HalM2 and HalM2:HalA2 complex were then imported into CIUSuite2 for analysis.[7-8] The replicate CIU data sets were averaged into a single data set and analyzed by the “Feature Detection” and “CIU50” modules of CIUSuite2. For feature detection, we used standard detection mode with a minimum feature length of 5 steps (20 V) and an allowed width in the drift time dimension of 0.3 ms. CIU50 was then employed with the default settings to fit the energy-resolved drift time profiles to estimate the collision voltage at which half of the protein had unfolded (Figure 4).

### Supporting Results

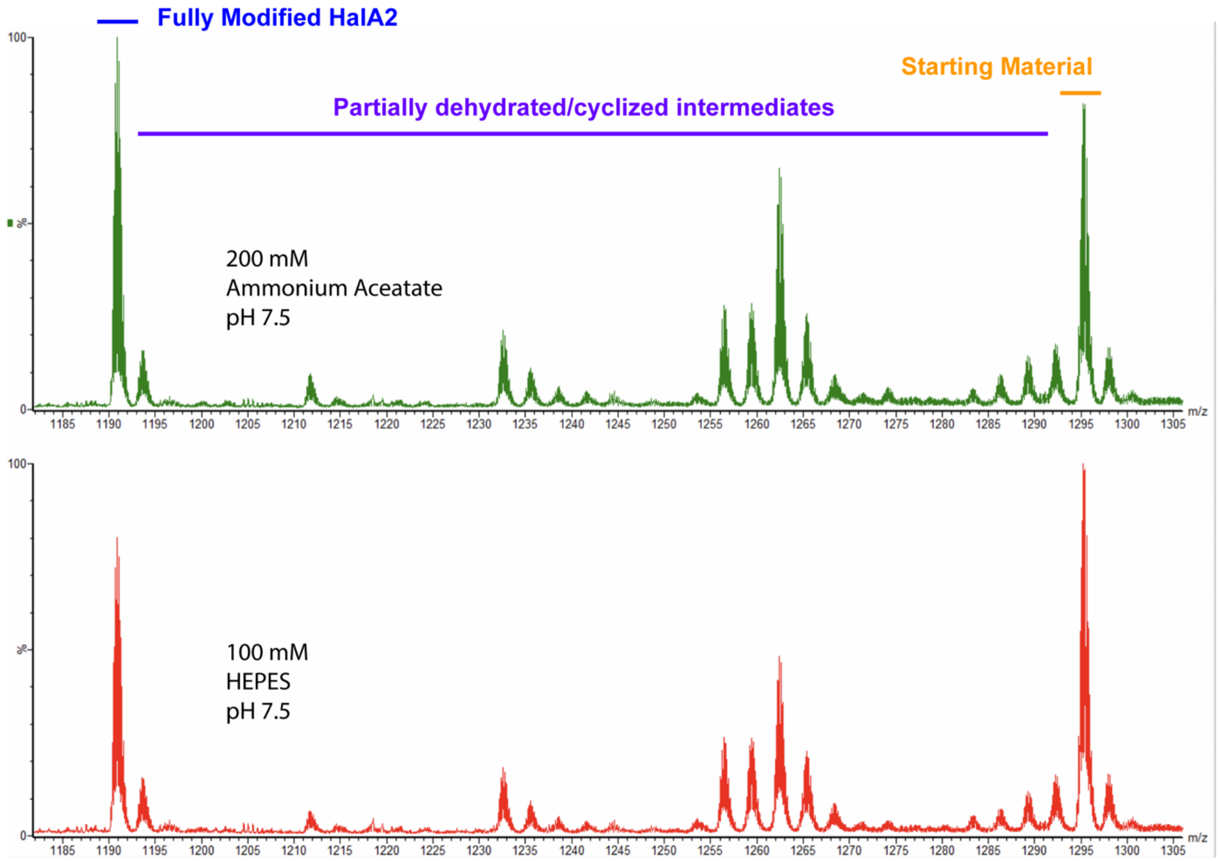


**Figure S1**. Comparison of HalM2 activity in the 200 mM ammonium acetate solvent used for native MS experiments (top panel) and the HEPES buffer used previously for kinetic studies of HalM2 (bottom panel).[1] Reactions were conducted for 5 min as described previously.[1] Reaction products were alkylated with *N*-ethylmaleimide to reveal the presence of thioether rings. The $7^+$ HalA2 ions are shown. The fully (4-fold) NEM alkylated starting material (containing no thioether rings) is indicated, along with the fully cyclized (unalkylated), 7-fold dehydrated final product. The other species correspond to partially cyclized and/or dehydrated intermediates. The activity in both solvents is nearly identical.

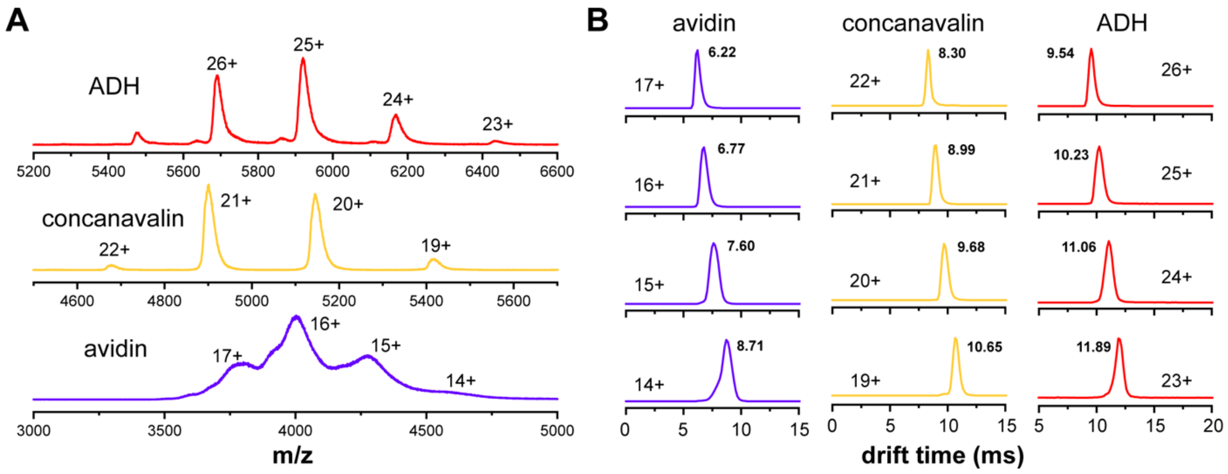


**Figure S2**. Native mass spectra (panel A) and ion mobiligrams (panel B) for the standard proteins used to generate the CCS calibration curve. Data were collected using the instrumental settings provided in Table S5. These data were used to construct the calibration curve (Figure S3) used for determination of the $\Omega_c$ and $\Omega_o$ values for HalM2, its variant enzymes, and their complexes with HalA2 (Tables 2, 3, S2 and S4).

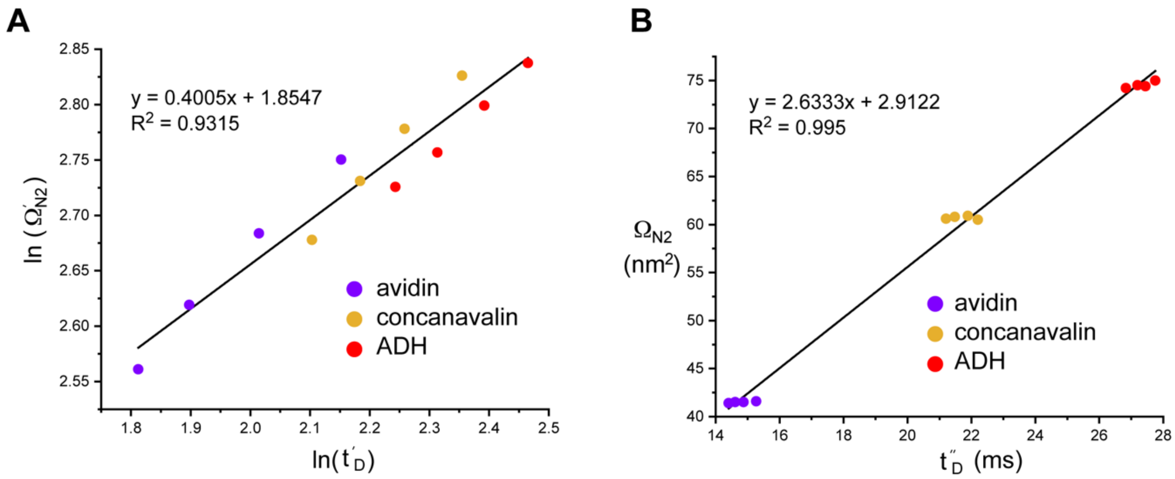


**Figure S3**. Calibration curves used to determine nitrogen collisional cross sections for HalM2, its variant enzymes, and their non-covalent complexes with HalA2. See Supporting Methods for details.

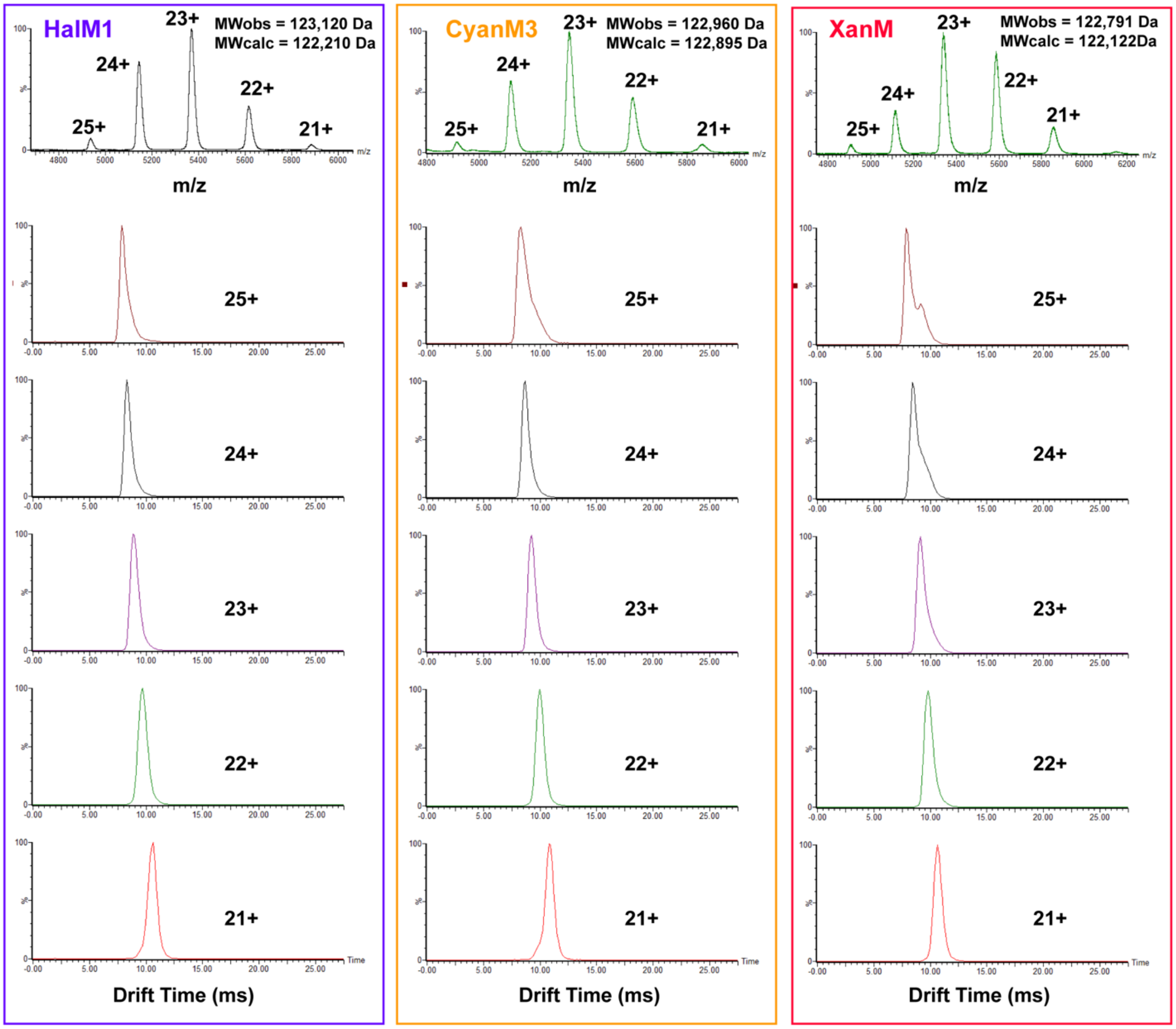


**Figure S4**. Native nESI-MS mass spectra and ion mobility arrival time distributions of other LanM enzymes. Data are shown for HalM1 from *Bacillus halodurans*,[9] CyanM3 from *Cyanothece sp.* PCC 7425,[10] and XanM from *Myxococcus xanthus*. The drift time distributions were fitted with sums of Gaussian peaks (Table S3) to determine the collisional cross sections of relevant conformations (Table S4). Each enzyme showed evidence for multiple conformations.

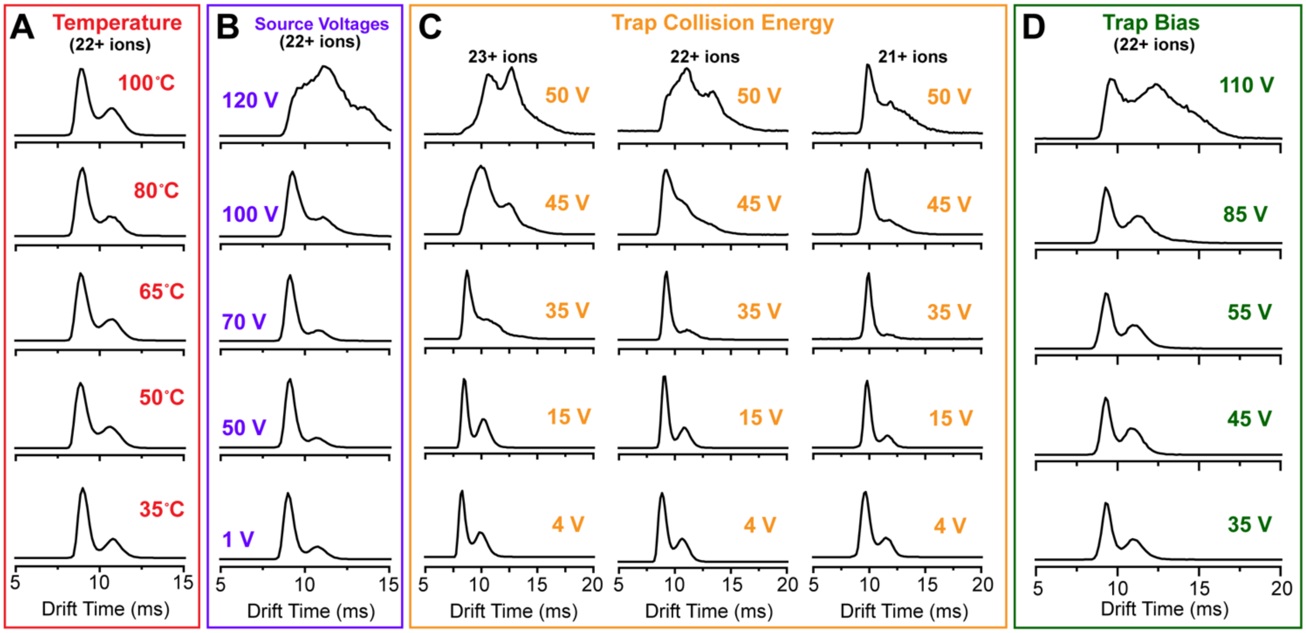


**Figure S5**. Control IM-MS experiments establish the stability of the gas phase HalM2 conformations over a large range of gentle ionization conditions. Ion mobility separations of the indicated HalM2 ions are shown over a range of source temperatures (panel A), cone and source offset voltages (panel B), trap collision energies (panel C), and trap bias voltages (panel D). The HalM2 conformational distribution was relatively insensitive to temperature, source voltages, and trap bias voltage (at a helium pressure of 5.2 mbar) over a large range of settings. Collisional activation performed in the trap at an argon pressure of 0.0245 mbar (panel C) resulted in unfolding of the open conformation near a trap collision energy of 30 V (a kinetic energy of 630-690 eV, depending on the HalM2 charge state) prior to unfolding of the compact conformation beginning around 40 V (840-920 eV). The instrumental settings for collection of these data are given in Table S5.

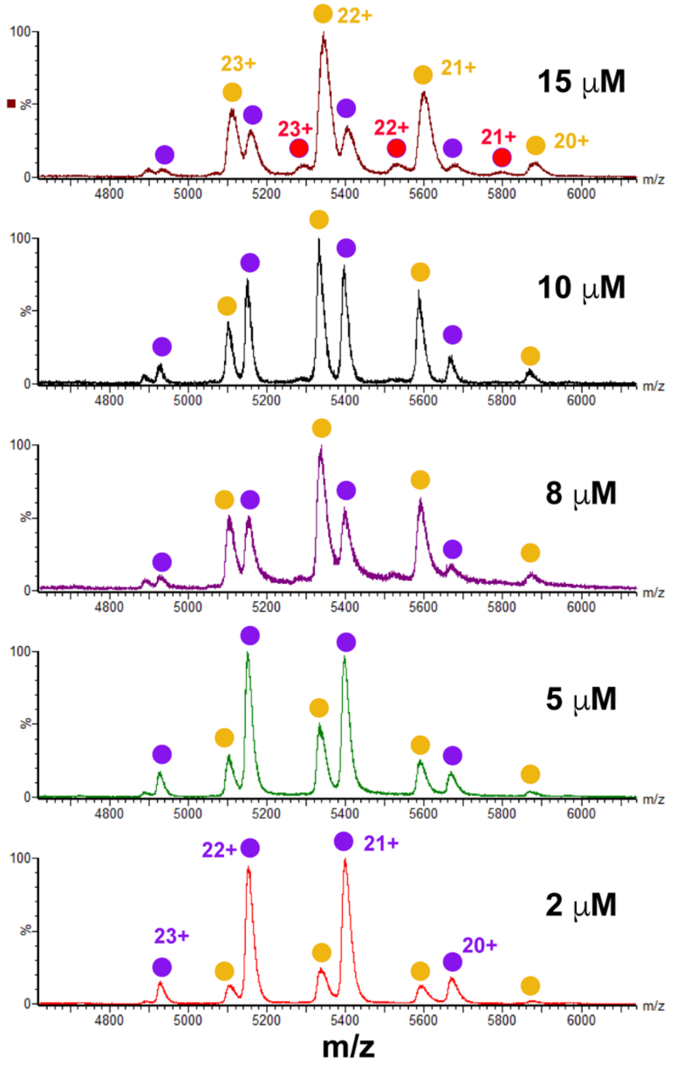


**Figure S6**. Native nESI mass spectra of the HalM2:HalA2 leader peptide complex. HalM2 (5 μM, purple circles) was incubated with the indicated concentration of the HalA2 leader peptide in 200 mM ammonium acetate and was electrosprayed using the instrumental conditions given in Table S5. The HalM2:HalA2 leader peptide complex (yellow circles) saturated as a function of HalA2 leader peptide concentration. At higher concentrations, a second equivalent of leader peptide bound to the enzymes (red circles).

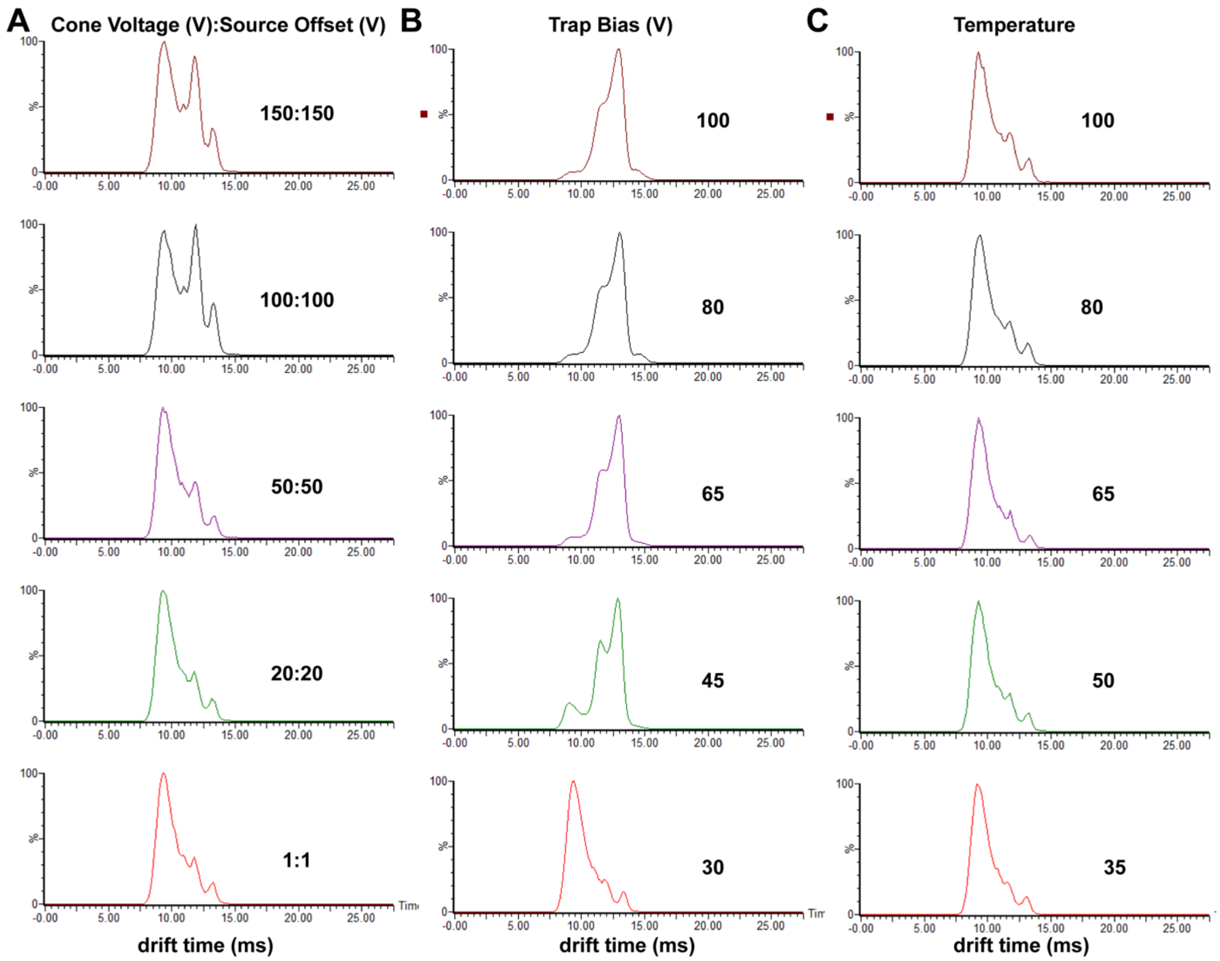


**Figure S7**. Dependence of HalA2 ion mobility arrival time distributions on selected instrumental parameters. Drift time distributions for the $[HalA2+5H]^{5+}$ ions are shown as a function of cone voltage and source offset voltage (panel A), trap bias voltage (panel B), and source temperature (panel C). Instrumental setting were otherwise as shown in Table S7.

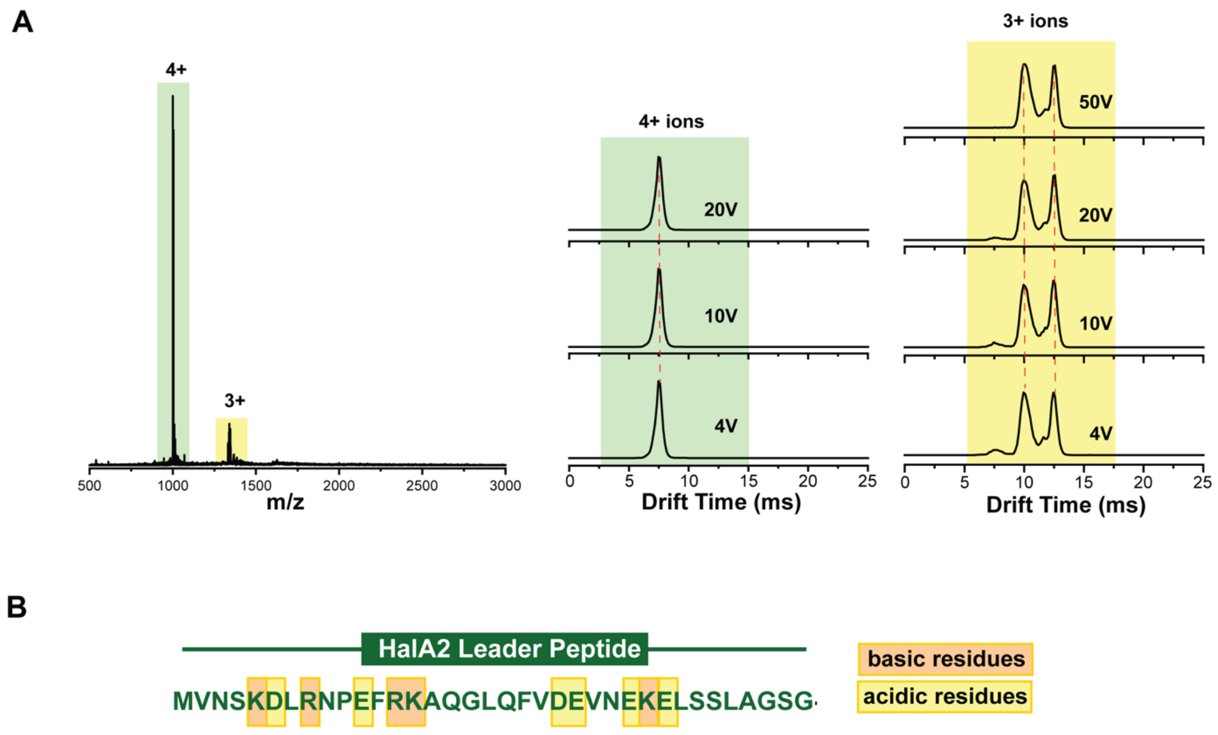


**Figure S8**. Native nESI-MS and ion mobility studies of the HalA2 leader peptide (HalA2 residues Met1-Gly36, $M_{obs} = 3992.89\ Da, M_{calc} = 3992.01\ Da$). A) Native mass spectrum (left) and ion mobility arrival time distributions (right) are shown for the $3^{+}$ and $4^{+}$ ions at the indicated trap collision energies. The $3^{+}$ ion shows some evidence for gas phase structure, perhaps involving formation of salt bridges between the acidic and basic residues in the leader peptide. Data were collected with 5 μM HalA2-LP using the instrumental settings given in Table S7.

**Table S1.** Determination of Molecular Weight ($MW$) for HalM2 and the HalM2:HalA2 complex.

| Protein | $Z$ | $(m/z)_{obs}$ | $A_{rel}$ | $MW$ |
|---|---|---|---|---|
| HalM2 | $19^+$ | $5963 \pm 1$ | $0.011 \pm 0.001$ | $113{,}287 \pm 25\ Da$ |
| | $20^+$ | $5565 \pm 1$ | $0.115 \pm 0.003$ | |
| | $21^+$ | $5395 \pm 1$ | $0.443 \pm 0.001$ | |
| | $22^+$ | $5149 \pm 1$ | $0.373 \pm 0.005$ | |
| | $23^+$ | $4924 \pm 1$ | $0.058 \pm 0.001$ | |
| HalM2:HalA2 complex | $20^+$ | $6028 \pm 3$ | $0.019 \pm 0.0005$ | $120{,}549\ \pm 20\ Da$ |
| | $21^+$ | $5740 \pm 1$ | $0.159 \pm 0.004$ | |
| | $22^+$ | $5479 \pm 1$ | $0.443 \pm 0.002$ | |
| | $23^+$ | $5241 \pm 1$ | $0.326 \pm 0.002$ | |
| | $24^+$ | $5022 \pm 1$ | $0.0525 \pm 0.0004$ | |

All values were determined in triplicate from the mass spectra of three separate samples. The standard error of the mean is reported for each parameter. The relative mass spectral peak area for each ion ($A_{rel}$) was used to calculate a weighted average for molecular weight ($MW$) for each individual spectrum from $(m/z)_{obs}$ and $Z$, respectively. These values were then averaged together to give the reported values. Data were collected with the instrumental settings provided in Table S5.

**Table S2**. Relevant observables for standard proteins used to determine collisional cross sections for lanthipeptide synthetases and their non-covalent complexes with precursor peptide.

| protein | $z^a$ | $(m/z)^a$ | $(t_D)^a$ | $(\Omega_{N2})^b$ | $(\Omega'_{N2})^c$ | $(t'_D)^d$ | $(t''_D)^e$ |
|---|---|---|---|---|---|---|---|
| avidin | $14^+$ | 4600 | 8.71 | 41.4 | 15.6 | 8.60 | 14.4 |
| | $15^+$ | 4278 | 7.6 | 41.5 | 14.6 | 7.50 | 14.6 |
| | $16^+$ | 4002 | 6.77 | 41.5 | 13.7 | 6.67 | 14.9 |
| | $17^+$ | 3785 | 6.22 | 41.6 | 12.9 | 6.12 | 15.3 |
| concanavalin | $19^+$ | 5417 | 10.65 | 60.6 | 16.9 | 10.53 | 21.2 |
| | $20^+$ | 5145 | 9.68 | 60.8 | 16.1 | 9.57 | 21.5 |
| | $21^+$ | 4899 | 8.99 | 60.9 | 15.3 | 8.88 | 21.9 |
| | $22^+$ | 4680 | 8.3 | 60.5 | 14.6 | 8.19 | 22.2 |
| ADH | $23^+$ | 6435 | 11.89 | 74.2 | 17.1 | 11.8 | 26.8 |
| | $24^+$ | 6169 | 11.06 | 74.5 | 16.4 | 10.94 | 27.2 |
| | $25^+$ | 5920 | 10.23 | 74.4 | 15.7 | 10.11 | 27.4 |
| | $26^+$ | 5692 | 9.54 | 75 | 15.3 | 9.42 | 27.8 |

[a]Measured from native mass spectra in Figure S2A
[b]Taken from Bush et. al. Anal. Chem. 2010, 82, 9557-9565
[c]Calculated from Equation 4
[d]Calculated from Equation 3
[e]Calculated from Equation 5
All drift times are in units of ms. All collisional cross section (Ω) values are in units of $nm^2$.

**Table S3**. Gaussian fits of ion mobility drift time distributions for wt HalM2, its variant enzymes, and their complexes with HalA2. Data were fitted with of Equation 1.

| Protein | $Z$ | Compact conformation | | | Open conformation | | | $y_0$ | $R^2$ |
|---|---|---|---|---|---|---|---|---|---|
| | | $x_c$ | $w$ | $A$ | $x_c$ | $w$ | $A$ | | |
| HalM2 | 20+ | 10.74 | 0.632 | 0.762 | 12.32 | 1.318 | 0.238 | -3.23 x 10-4 | 0.99 |
| | 21+ | 9.87 | 0.580 | 0.684 | 11.48 | 1.248 | 0.316 | -6.57 x 10-4 | 0.99 |
| | 22+ | 9.19 | 0.499 | 0.603 | 10.73 | 1.230 | 0.397 | -2.86 x 10-4 | 0.99 |
| | 23+ | 8.64 | 0.492 | 0.507 | 10.12 | 1.284 | 0.493 | 2.48 x 10-4 | 0.99 |
| HalM2:HalA2 complex | 21+ | 10.90 | 0.723 | 0.709 | 11.71 | 1.320 | 0.291 | 3.24 x 10-4 | 0.99 |
| | 22+ | 10.13 | 0.645 | 0.573 | 11.00 | 1.219 | 0.427 | 1.53 x 10-4 | 0.99 |
| | 23+ | 9.42 | 0.524 | 0.389 | 10.29 | 1.214 | 0.611 | 1.55 x 10-4 | 0.99 |
| | 24+ | 8.81 | 0.492 | 0.264 | 9.78 | 1.262 | 0.736 | 0.00275 | 0.98 |
| HalM2:HalA2-LP complex | 21+ | 10.36 | 0.592 | 0.713 | 11.37 | 1.569 | 0.287 | 6.13 x 10-4 | 0.99 |
| | 22+ | 9.63 | 0.578 | 0.593 | 10.96 | 1.268 | 0.407 | -1.51 x 10-4 | 0.99 |
| | 23+ | 9.04 | 0.568 | 0.485 | 10.41 | 1.054 | 0.515 | 6.52 x 10-4 | 0.99 |
| | 24+ | 8.53 | 0.584 | 0.369 | 9.88 | 1.176 | 0.631 | 0.00989 | 0.98 |
| GS491 | 20+ | 10.70 | 0.579 | 0.727 | 11.82 | 1.728 | 0.273 | 0.00119 | 0.99 |
| | 21+ | 9.869 | 0.543 | 0.676 | 11.14 | 1.520 | 0.324 | -1.58 x 10-4 | 0.99 |
| | 22+ | 9.201 | 0.482 | 0.613 | 10.44 | 1.439 | 0.387 | -3.524 x 10-4 | 0.99 |
| | 23+ | 8.666 | 0.486 | 0.555 | 9.93 | 1.373 | 0.445 | 0.00131 | 0.99 |
| GS491:HalA2 complex | 21+ | 10.78 | 0.712 | 0.747 | 11.56 | 1.311 | 0.253 | 0.0024 | 0.99 |
| | 22+ | 10.05 | 0.646 | 0.593 | 10.92 | 1.277 | 0.407 | 0.00121 | 0.99 |
| | 23+ | 9.378 | 0.527 | 0.394 | 10.24 | 1.234 | 0.606 | 8.7 x 0-4 | 0.99 |
| | 24+ | 8.796 | 0.496 | 0.325 | 9.777 | 1.257 | 0.675 | 0.00744 | 0.98 |
| GS116 | 20+ | 10.77 | 0.580 | 0.786 | 11.70 | 1.612 | 0.214 | 0.00185 | 0.99 |
| | 21+ | 11.22 | 1.595 | 0.302 | 9.951 | 0.556 | 0.698 | 1.351 x 10-4 | 0.99 |
| | 22+ | 9.268 | 0.521 | 0.627 | 10.65 | 1.377 | 0.373 | -3.309 x 10-4 | 0.99 |
| | 23+ | 8.697 | 0.526 | 0.588 | 10.08 | 1.278 | 0.412 | 3.884 x 10-4 | 0.99 |
| GS116: HalA2 complex | 21+ | 10.88 | 0.724 | 0.731 | 11.74 | 1.339 | 0.269 | 0.00408 | 0.99 |
| | 22+ | 10.12 | 0.650 | 0.569 | 11.03 | 1.316 | 0.431 | 0.00194 | 0.99 |
| | 23+ | 9.428 | 0.559 | 0.411 | 10.36 | 1.302 | 0.589 | 0.00146 | 0.99 |
| | 24+ | 8.798 | 0.565 | 0.402 | 9.929 | 1.199 | 0.598 | 0.00746 | 0.97 |
| GS389 | 19+ | 11.65 | 0.737 | 1.000 | - | - | | 0.00784 | 0.98 |
| | 20+ | 10.61 | 0.571 | 0.699 | 11.11 | 0.968 | 0.301 | 0.00282 | 0.99 |
| | 21+ | 9.832 | 0.539 | 0.664 | 10.67 | 1.396 | 0.336 | 7.69 x 10-4 | 0.99 |
| | 22+ | 9.185 | 0.511 | 0.533 | 10.26 | 1.449 | 0.467 | 4.9 x 10-4 | 0.99 |
| GS389: HalA2 complex | 20+ | 11.50 | 0.677 | 1.000 | - | - | | 0.00486 | 0.99 |
| | 21+ | 10.54 | 0.569 | 0.673 | 11.16 | 1.091 | 0.327 | 0.00259 | 0.99 |
| | 22+ | 9.84 | 0.551 | 0.509 | 10.67 | 1.283 | 0.491 | 0.00147 | 0.99 |
| | 23+ | 9.24 | 0.489 | 0.326 | 10.10 | 1.241 | 0.674 | 0.00135 | 0.99 |
| GS635 | 19+ | 11.58 | 0.563 | 0.348 | 12.06 | 1.083 | 0.652 | 0.00218 | 0.99 |
| | 20+ | 10.67 | 0.601 | 0.347 | 11.28 | 1.081 | 0.653 | 8.233 x 10-4 | 0.99 |
| | 21+ | 9.909 | 0.535 | 0.298 | 10.53 | 1.058 | 0.702 | 7.616 x 10-4 | 0.99 |

| | | | | | | | | | |
|---|---|---|---|---|---|---|---|---|---|
| | 22+ | 9.234 | 0.439 | 0.173 | 9.916 | 1.133 | 0.827 | 0.00245 | 0.99 |
| GS635:HalA2 complex | 20+ | 11.57 | 0.636 | 0.416 | 12.04 | 0.997 | 0.584 | 0.00162 | 0.99 |
| | 21+ | 10.67 | 0.602 | 0.385 | 11.20 | 0.985 | 0.615 | 8.52 x $10^{-4}$ | 0.99 |
| | 22+ | 9.993 | 0.592 | 0.391 | 10.55 | 0.988 | 0.609 | 6.114 x $10^{-4}$ | 0.99 |
| | 23+ | 9.444 | 0.536 | 0.204 | 9.972 | 1.008 | 0.796 | 0.0019 | 0.99 |
| GS341 | 19+ | 11.71 | 0.671 | 0.357 | 12.08 | 1.000 | 0.643 | 0.00397 | 0.99 |
| | 20+ | 10.82 | 0.672 | 0.450 | 11.32 | 0.976 | 0.550 | 0.00137 | 0.99 |
| | 21+ | 10.12 | 0.722 | 0.502 | 10.64 | 0.999 | 0.498 | 0.00107 | 0.99 |
| | 22+ | 9.687 | 0.864 | 0.586 | 10.27 | 10.158 | 0.414 | 0.00554 | 0.97 |
| GS341:HalA2 complex | 20+ | 11.66 | 0.670 | 0.513 | 12.12 | 0.940 | 0.487 | 0.00212 | 0.99 |
| | 21+ | 10.73 | 0.600 | 0.421 | 11.22 | 0.952 | 0.579 | 0.00142 | 0.99 |
| | 22+ | 10.06 | 0.618 | 0.483 | 10.64 | 1.015 | 0.517 | 0.00118 | 0.99 |
| | 23+ | 9.449 | 0.562 | 0.222 | 10.04 | 1.073 | 0.778 | 0.00558 | 0.98 |
| GS331 | 19+ | 11.70 | 0.677 | 0.351 | 11.88 | 1.098 | 0.649 | 0.00199 | 0.99 |
| | 20+ | 10.71 | 0.681 | 0.503 | 11.19 | 0.957 | 0.497 | 5.061 x $10^{-4}$ | 0.99 |
| | 21+ | 9.971 | 0.672 | 0.505 | 10.50 | 0.970 | 0.495 | 2.22 x $10^{-4}$ | 0.99 |
| | 22+ | 9.485 | 0.786 | 0.584 | 10.01 | 1.057 | 0.416 | 9.041 x $10^{-4}$ | 0.99 |
| GS331:HalA2 complex | 20+ | 11.53 | 0.657 | 0.456 | 11.96 | 0.947 | 0.544 | 0.00114 | 0.99 |
| | 21+ | 10.63 | 0.622 | 0.439 | 11.13 | 0.948 | 0.561 | 4.608 x $10^{-4}$ | 0.99 |
| | 22+ | 9.920 | 0.586 | 0.431 | 10.46 | 0.964 | 0.569 | 3.631 x $10^{-4}$ | 0.99 |
| | 23+ | 9.338 | 0.524 | 0.228 | 9.872 | 0.986 | 0.772 | 0.00108 | 0.99 |
| HalM1 | 21+ | 10.56 | 0.773 | 1.000 | - | - | | 330 | 0.99 |
| | 22+ | 9.71 | 0.777 | 1.000 | - | - | | 1606 | 0.99 |
| | 23+ | 8.87 | 0.533 | 0.592 | 9.31 | 0.826 | 0.408 | 1442 | 0.98 |
| | 24+ | 8.31 | 0.451 | 0.605 | 8.77 | 0.831 | 0.395 | 1037 | 0.99 |
| | 25+ | 7.88 | 0.426 | 0.509 | 8.39 | 0.920 | 0.491 | 281 | 0.99 |
| CyanM3 | 21+ | 10.84 | 0.760 | 1.000 | - | - | | 1375 | 0.99 |
| | 22+ | 10.00 | 0.705 | 1.000 | - | - | | 3467 | 0.99 |
| | 23+ | 9.19 | 0.526 | 0.651 | 9.60 | 0.783 | 0.349 | 2038 | 0.99 |
| | 24+ | 8.64 | 0.476 | 0.612 | 9.12 | 0.867 | 0.388 | 1465 | 0.99 |
| | 25+ | 8.30 | 0.652 | 0.507 | 9.15 | 1.422 | 0.493 | 122 | 0.99 |
| XanM | 21+ | 10.71 | 0.719 | 1.000 | - | - | | 10883 | 0.99 |
| | 22+ | 9.84 | 0.617 | 0.689 | 10.47 | 1.176 | 0.311 | 19106 | 0.99 |
| | 23+ | 9.16 | 0.566 | 0.531 | 9.87 | 1.286 | 0.469 | 17283 | 0.99 |
| | 24+ | 8.51 | 0.524 | 0.445 | 9.38 | 1.396 | 0.555 | 9470 | 0.99 |
| | 25+ | 7.97 | 0.527 | 0.455 | 9.06 | 1.455 | 0.545 | 3292 | 0.99 |

Table S4. Collisional cross sections of wt HalM2, its variant enzymes, and their complexes with HalA2.

| Protein or Complex | Z | Compact Conformation Drift Time (ms) | $\Omega_c$ $(nm^2)$ | $\Omega_{c,avg}$ $(nm^2)$ | Open Conformation Drift Time (ms) | $\Omega_o$ $(nm^2)$ | $\Omega_{o,avg}$ $(nm^2)$ | $\Delta\Omega_{free}$ or $\Delta\Omega_{complex}$ |
|---|---|---|---|---|---|---|---|---|
| HalM2 | 20+ | $10.74 \pm 0.002$ | 61.6 | $63.1 \pm 0.7$ | $12.32 \pm 0.02$ | 64.9 | $66.9 \pm 0.8$ | 3.8 |
| | 21+ | $9.87 \pm 0.002$ | 62.4 | | $11.48 \pm 0.01$ | 66.2 | | |
| | 22+ | $9.19 \pm 0.002$ | 63.5 | | $10.73 \pm 0.01$ | 67.5 | | |
| | 23+ | $8.64 \pm 0.002$ | 64.7 | | $10.12 \pm 0.01$ | 68.8 | | |
| HalM2:HalA2 complex | 21+ | $10.89 \pm 0.004$ | 64.9 | $66.4 \pm 0.7$ | $11.71 \pm 0.09$ | 66.7 | $68.7 \pm 0.9$ | 2.3 |
| | 22+ | $10.13 \pm 0.003$ | 66.0 | | $11.00 \pm 0.03$ | 68.1 | | |
| | 23+ | $9.42 \pm 0.003$ | 66.9 | | $10.29 \pm 0.02$ | 69.3 | | |
| | 24+ | $8.81 \pm 0.005$ | 67.9 | | $9.78 \pm 0.01$ | 70.8 | | |
| HalM2:HalA2-LP complex | 21+ | $10.35 \pm 0.002$ | 63.6 | $65.3 \pm 0.8$ | $11.37 \pm 0.04$ | 65.9 | $68.6 \pm 1.0$ | 3.3 |
| | 22+ | $9.63 \pm 0.002$ | 64.7 | | $10.96 \pm 0.01$ | 68.0 | | |
| | 23+ | $9.04 \pm 0.002$ | 65.9 | | $10.41 \pm 0.004$ | 69.6 | | |
| | 24+ | $8.53 \pm 0.005$ | 67.1 | | $9.88 \pm 0.01$ | 71.0 | | |
| GS491 | 20+ | $10.69 \pm 0.002$ | 61.4 | $63.1 \pm 0.7$ | $11.82 \pm 0.06$ | 63.9 | $66.1 \pm 0.9$ | 3.0 |
| | 21+ | $9.87 \pm 0.002$ | 62.4 | | $11.14 \pm 0.03$ | 65.4 | | |
| | 22+ | $9.2 \pm 0.002$ | 63.5 | | $10.44 \pm 0.02$ | 66.7 | | |
| | 23+ | $8.66 \pm 0.002$ | 64.8 | | $9.93 \pm 0.01$ | 68.3 | | |
| GS491:HalA2 complex | 21+ | $10.78 \pm 0.003$ | 64.6 | $66.3 \pm 0.7$ | $11.5 \pm 0.1$ | 66.4 | $68.5 \pm 0.9$ | 2.2 |
| | 22+ | $10.05 \pm 0.003$ | 65.8 | | $10.91 \pm 0.03$ | 67.9 | | |
| | 23+ | $9.38 \pm 0.002$ | 66.8 | | $10.24 \pm 0.01$ | 69.1 | | |
| | 24+ | $8.79 \pm 0.004$ | 67.9 | | $9.78 \pm 0.01$ | 70.8 | | |
| GS116 | 20+ | $10.77 \pm 0.002$ | 61.6 | $63.2 \pm 0.7$ | $11.69 \pm 0.07$ | 63.6 | $66.3 \pm 1.1$ | 3.1 |
| | 21+ | $9.95 \pm 0.04$ | 62.6 | | $11.22 \pm 0.002$ | 65.6 | | |
| | 22+ | $9.27 \pm 0.002$ | 63.7 | | $10.65 \pm 0.01$ | 67.3 | | |
| | 23+ | $8.69 \pm 0.002$ | 64.9 | | $10.08 \pm 0.01$ | 68.7 | | |
| GS116:HalA2 complex | 21+ | $10.88 \pm 0.003$ | 64.8 | $66.4 \pm 0.7$ | $11.73 \pm 0.07$ | 66.8 | $68.9 \pm 0.9$ | 2.5 |
| | 22+ | $10.12 \pm 0.002$ | 65.9 | | $11.03 \pm 0.03$ | 68.2 | | |
| | 23+ | $9.43 \pm 0.002$ | 67.0 | | $10.36 \pm 0.01$ | 69.5 | | |
| | 24+ | $8.79 \pm 0.006$ | 67.9 | | $9.93 \pm 0.01$ | 71.2 | | |
| GS389 | 19+ | $11.65 \pm 0.002$ | 60.5 | $61.9 \pm 0.7$ | - | - | $64.3 \pm 1.1$ | 2.4 |
| | 20+ | $10.61 \pm 0.003$ | 61.3 | | $11.11 \pm 0.09$ | 62.4 | | |
| | 21+ | $9.83 \pm 0.002$ | 62.3 | | $10.67 \pm 0.05$ | 64.4 | | |

| | | | | | | | | |
|---|---|---|---|---|---|---|---|---|
| | $22^+$ | $9.19 \pm 0.002$ | 63.5 | | $10.26 \pm 0.02$ | 66.3 | | |
| GS389:HalA2 complex | $20^+$ | $11.50 \pm 0.002$ | 63.2 | $64.7 \pm 0.7$ | - | - | $67.2 \pm 1.0$ | 2.5 |
| | $21^+$ | $10.53 \pm 0.002$ | 64.0 | | $11.16 \pm 0.06$ | 65.4 | | |
| | $22^+$ | $9.84 \pm 0.002$ | 65.2 | | $10.67 \pm 0.03$ | 67.3 | | |
| | $23^+$ | $9.23 \pm 0.002$ | 66.4 | | $10.10 \pm 0.01$ | 68.6 | | |
| GS635 | $19^+$ | $11.58 \pm 0.003$ | 60.3 | $62.0 \pm 0.7$ | $12.06 \pm 0.01$ | 61.3 | $63.4 \pm 0.9$ | 1.4 |
| | $20^+$ | $10.67 \pm 0.004$ | 61.4 | | $11.28 \pm 0.02$ | 62.7 | | |
| | $21^+$ | $9.91 \pm 0.003$ | 62.5 | | $10.53 \pm 0.02$ | 64.0 | | |
| | $22^+$ | $9.23 \pm 0.004$ | 63.6 | | $9.92 \pm 0.01$ | 65.4 | | |
| GS635:HalA2 complex | $20^+$ | $11.57 \pm 0.006$ | 63.3 | $65.1 \pm 0.8$ | $12.04 \pm 0.05$ | 64.3 | $66.3 \pm 0.9$ | 1.2 |
| | $21^+$ | $10.67 \pm 0.006$ | 64.3 | | $11.20 \pm 0.04$ | 65.6 | | |
| | $22^+$ | $9.99 \pm 0.005$ | 65.6 | | $10.55 \pm 0.04$ | 67.0 | | |
| | $23^+$ | $9.44 \pm 0.006$ | 67.0 | | $9.97 \pm 0.02$ | 68.4 | | |
| GS341 | $19^+$ | $11.71 \pm 0.01$ | 60.6 | $62.5 \pm 0.9$ | $12.07 \pm 0.04$ | 61.3 | $63.7 \pm 1.1$ | 1.2 |
| | $20^+$ | $10.82 \pm 0.01$ | 61.7 | | $11.3 \pm 0.1$ | 62.8 | | |
| | $21^+$ | $10.12 \pm 0.03$ | 63.0 | | $10.6 \pm 0.2$ | 64.3 | | |
| | $22^+$ | $9.68 \pm 0.08$ | 64.8 | | $10.3 \pm 0.7$ | 66.3 | | |
| GS341:HalA2 complex | $20^+$ | $11.66 \pm 0.02$ | 63.5 | $65.2 \pm 0.8$ | $12.1 \pm 0.1$ | 64.5 | $66.5 \pm 0.9$ | 1.3 |
| | $21^+$ | $10.73 \pm 0.008$ | 64.5 | | $11.22 \pm 0.06$ | 65.6 | | |
| | $22^+$ | $10.06 \pm 0.007$ | 65.8 | | $10.64 \pm 0.07$ | 67.2 | | |
| | $23^+$ | $9.45 \pm 0.01$ | 67.0 | | $10.03 \pm 0.04$ | 68.6 | | |
| GS331 | $19^+$ | $11.69 \pm 0.009$ | 60.5 | $62.3 \pm 0.8$ | $11.88 \pm 0.02$ | 60.9 | $63.3 \pm 1.0$ | 1.0 |
| | $20^+$ | $10.71 \pm 0.009$ | 61.5 | | $11.19 \pm 0.08$ | 62.5 | | |
| | $21^+$ | $9.97 \pm 0.01$ | 62.7 | | $10.49 \pm 0.07$ | 63.9 | | |
| | $22^+$ | $9.49 \pm 0.02$ | 64.3 | | $10.01 \pm 0.2$ | 65.7 | | |
| GS331:HalA2 complex | $20^+$ | $11.53 \pm 0.006$ | 63.3 | $64.9 \pm 0.7$ | $11.97 \pm 0.05$ | 64.2 | $66.1 \pm 0.9$ | 1.2 |
| | $21^+$ | $10.69 \pm 0.007$ | 64.4 | | $11.13 \pm 0.05$ | 65.4 | | |
| | $22^+$ | $9.92 \pm 0.004$ | 65.4 | | $10.46 \pm 0.04$ | 66.8 | | |
| | $23^+$ | $9.34 \pm 0.004$ | 66.7 | | $9.87 \pm 0.01$ | 68.2 | | |
| HalM1 | $21^+$ | $10.56 \pm 0.001$ | 64.1 | $65.7 \pm 0.6$ | - | - | $67.9 \pm 0.8$ | 2.2 |
| | $22^+$ | $9.71 \pm 0.002$ | 64.9 | | - | - | | |
| | $23^+$ | $8.87 \pm 0.01$ | 65.4 | | $9.3 \pm 0.1$ | 66.6 | | |
| | $24^+$ | $8.31 \pm 0.003$ | 66.4 | | $8.77 \pm 0.05$ | 67.8 | | |
| | $25^+$ | $7.88 \pm 0.002$ | 67.7 | | $8.39 \pm 0.03$ | 69.3 | | |
| | $21^+$ | $10.84 \pm 0.003$ | 64.7 | | - | - | | |

| | | | | | | | | |
|---|---|---|---|---|---|---|---|---|
| CyanM3 | $22^+$ | $10.00 \pm 0.002$ | 65.6 | $66.0 \pm 0.7$ | - | - | $69.3 \pm 1.3$ | 3.3 |
| | $23^+$ | $9.19 \pm 0.01$ | 66.3 | | $9.6 \pm 0.1$ | 67.4 | | |
| | $24^+$ | $8.64 \pm 0.002$ | 67.4 | | $9.12 \pm 0.05$ | 68.9 | | |
| | $25^+$ | $8.30 \pm 0.01$ | 69.1 | | $9.15 \pm 0.07$ | 71.7 | | |
| XanM | $21^+$ | $10.71 \pm 0.004$ | 64.4 | $66.1 \pm 0.6$ | - | - | $69.0 \pm 1.0$ | 2.9 |
| | $22^+$ | $9.84 \pm 0.004$ | 65.21 | | $10.5 \pm 0.1$ | 66.8 | | |
| | $23^+$ | $9.16 \pm 0.003$ | 66.1 | | $9.87 \pm 0.04$ | 68.2 | | |
| | $24^+$ | $8.51 \pm 0.003$ | 67.0 | | $9.38 \pm 0.03$ | 69.6 | | |
| | $25^+$ | $7.97 \pm 0.004$ | 68.0 | | $9.06 \pm 0.03$ | 71.5 | | |

Drift times are in units of ms. Collisional cross sections (Ω) are in units of $nm^2$. All drift times were determined by fitting the combined data for three replicates to a sum of two Gaussian-shaped peaks (Table S3). The reported error on the drift time is the standard error provided by the Gaussian fit. Collisional cross sections were calculated from the drift times as described in the Supporting Information from standard curves (Figure S3) generated from a set of proteins with known $\Omega_{N2}$ values (Figure S2, Table S2). $\Omega_{avg} \pm s.e.m.$ were calculated as the mean of the collisional cross sections of the individual charge states. Data were collected with the instrumental settings provided in Table S5.

**Table S5**. Instrumental conditions used for determination of the molecular weight ($MW$) and collisional cross sections ($\Omega_c$ and $\Omega_o$) of HalM2, its variant enzymes, their non-covalent complexes with HalA2, and the CCS standard proteins (Table S1).

| **Prior to IM analysis** | |
|---|---|
| **setting** | **value** |
| Capillary voltage | 1.5 kV |
| Cone voltage | 1 V |
| Source offset | 1 V |
| Source temperature | 35 °C |
| Trap collision energy | 4 V |
| Trap gas flow (argon) | 2 mL/min |
| He cell gas flow | 180 mL/min |
| Trap DC Entrance | 3.0 V |
| Trap DC Bias | 35 V |
| Trap DC | 0 |
| Trap DC exit | 0 |
| Trap wave velocity | 311 m/s |
| Trap wave height | 4 V |
| **During IM analysis** | |
| **setting** | **value** |
| IMS gas flow (nitrogen) | 90 mL/min |
| IMS DC entrance | 20 V |
| Helium cell DC | 50 V |
| Helium exit | -20 V |
| IMS bias | 3.0 |
| IMS DC exit | 0 |
| IMS wave velocity | 550 m/s |
| IMS wave height | 40 V |
| Mobility trapping release time | 500 μs |
| Mobility trapping height | 15 V |
| Mobility extract height | 0 V |
| **Following IM analysis** | |
| **setting** | **value** |
| Transfer collision energy | OFF |
| Transfer DC entrance | 5 V |
| Transfer DC exit | 15 V |
| Transfer wave velocity | 175 m/s |
| Transfer wave height | 4 V |

**Table S6**. Instrumental conditions used for Collision Induced Unfolding (CIU) Studies of HalM2 and the HalM2:HalA2 complex

| **Prior to IM analysis** | |
|---|---|
| **setting** | **value** |
| Capillary voltage | 1.5 kV |
| Cone voltage | 50 V |
| Source offset | 50 V |
| Source temperature | 70 °C |
| Trap collision energy* | 5 V |
| Trap gas flow (argon) | 10 mL/min |
| He cell gas flow | 180 mL/min |
| Trap DC Entrance | 3.0 V |
| Trap DC Bias | 45 V |
| Trap DC | 0 |
| Trap DC exit | 0 |
| Trap wave velocity | 311 m/s |
| Trap wave height | 4 V |
| **During IM analysis** | |
| **setting** | **value** |
| IMS gas flow (nitrogen) | 90 mL/min |
| IMS DC entrance | 20 V |
| Helium cell DC | 50 V |
| Helium exit | -20 V |
| IMS bias | 3.0 |
| IMS DC exit | 0 |
| IMS wave velocity | 550 m/s |
| IMS wave height | 40 V |
| Mobility trapping release time | 500 μs |
| Mobility trapping height | 15 V |
| Mobility extract height | 0 V |
| **Following IM analysis** | |
| **setting** | **value** |
| Transfer collision energy | OFF |
| Transfer DC entrance | 5 V |
| Transfer DC exit | 15 V |
| Transfer wave velocity | 175 m/s |
| Transfer wave height | 4 V |

*Trap CE was varied from 5-180 V in 5 V increments

**Table S7**. Instrumental conditions used for Ion Mobility studies of HalA2 and the HalA2 leader peptide

| setting | value |
|---|---|
| **Prior to the analysis** | |
| **setting** | **value** |
| Capillary voltage | 1.5 kV |
| Cone voltage | 1 V |
| Source offset | 1 V |
| Source temperature | 70 °C |
| Trap collision energy | 4 V |
| Trap gas flow (argon) | 2 mL/min |
| He cell gas flow | 180 mL/min |
| Trap DC Entrance | 3.0 V |
| Trap DC Bias | 30 V |
| Trap DC | 0 |
| Trap DC exit | 0 |
| Trap wave velocity | 311 m/s |
| Trap wave height | 4 V |
| **During IM analysis** | |
| **setting** | **value** |
| IMS gas flow (nitrogen) | 90 mL/min |
| IMS DC entrance | 20 V |
| Helium cell DC | 50 V |
| Helium exit | -20 V |
| IMS bias | 3.0 |
| IMS DC exit | 0 |
| IMS wave velocity | 400 m/s |
| IMS wave height | 25 V |
| Mobility trapping release time | 500 μs |
| Mobility trapping height | 15 V |
| Mobility extract height | 0 V |
| **Following IM analysis** | |
| **setting** | **value** |
| Transfer collision energy | OFF |
| Transfer DC entrance | 5 V |
| Transfer DC exit | 15 V |
| Transfer wave velocity | 175 m/s |
| Transfer wave height | 4 V |
